# MGM2 as a Multimodal Foundation Model for Microbial Community States and Responses

**DOI:** 10.64898/2026.07.20.739063

**Authors:** Haohong Zhang, Yuli Zhang, Yuxuan Qi, Tianao Liu, Linfeng Zhao, Ronghua Yang, Kang Ning

## Abstract

Microbial communities sustain ecosystems and influence host health, yet connecting their molecular diversity to responses under intervention remains difficult. Here we introduce MGM2, a multimodal foundation model integrating DNA language-model representations of microbial sequences, quantitative abundances and sample descriptions. After pretraining on 1.8 million microbiomes, its frozen representations achieved a mean macro-AUROC of 0.91 on later MGnify projects absent from pretraining. These representations also supported fecal microbiota transplantation outcome prediction, recovering 68% of observed high-change taxa. Adding molecular structures enabled prediction of drug effects in nested five-fold evaluation, while measured perturbed states informed subsequent recovery. To interpret the learned representations, sparse decomposition resolved taxonomic, ecological and abundance-related features. Together, these results show how a shared representation of molecular identity and community context connects large-scale microbiome learning to the prediction and interpretation of community change.

**Highlights:**

- Multimodal pretraining integrates microbial sequence, abundance and biological descriptions to learn community representations that transfer across studies and ecological settings.
- Paired donor–recipient representations support prediction of clinical FMT response and post-transplant composition, including the taxa undergoing the largest abundance changes.
- Community states combined with molecular structure support prediction of drug effects on community composition, while post-perturbation states inform the direction of subsequent recovery.
- Sparse features distinguish microbial identity from ecological and abundance context, while molecular attention interventions expose drug-input sensitivity for biological follow-up.

## Introduction

Microbial communities sustain biogeochemical cycles and shape host metabolism, making their organization central to ecosystem function and human health [1, 2]. Long before microbiome sequencing became routine, molecular phylogeny showed that conserved sequences could reveal the deep evolutionary organization of microbial life [3, 4]. Large-scale surveys subsequently placed these molecular identities within communities distributed across human hosts and the biosphere [5, 6]. Resources such as MGnify, Qiita and MicrobeAtlas now connect observations across studies and habitats [7–9]. Yet identifying the members of a community is only part of understanding it: the same organisms encounter different neighbors, resources and disturbances in different settings [10]. Relating molecular identity to this ecological context is essential both for comparing microbiomes and for anticipating how they respond to introduced organisms or chemical compounds.

Biological foundation models offer a way to learn relationships that can be reused across these settings. Genomic language models, including Nucleotide Transformer and Evo 2, learn sequence representations across diverse organisms [11, 12], while ProkBERT and METAGENE-1 target microbial genomic and metagenomic applications [13, 14]. Community models instead learn from the organisms observed together: Pope et al. pretrained a microbiome language model on gut profiles [15], our previous Microbial General Model (MGM) learned across 260,000 samples from diverse contexts [16], and Waypoint extended taxonomic-profile modeling to ecological classification and drug-related endpoints [17]. Susagi brings sequence information into community modeling through sets of bacterial SSU rRNA gene embeddings, learning member-level stability without abundance inputs [18]. These approaches connect different levels of microbial biology. The next step is to integrate molecular relationships with quantitative community composition and ecological descriptions in a representation that can support both community analysis and intervention-response prediction.

The information needed for this connection spans several modalities. Nucleotide sequence provides a molecular reference for microbial identity, abundance records the quantitative structure of a community, and sample descriptions locate that community within a host, habitat or experimental setting. Abundance-based models have established the predictive value of composition [19, 20], although compositionality and variable feature definitions complicate comparisons across datasets [21]. Sequence representations can relate microbial features through molecular similarity, while descriptions supply context absent from a count matrix. Integrating these inputs would allow an organism to retain its molecular reference while its encoded state changes with abundance and neighboring organisms.

Predicting community responses adds a second requirement: linking the community state to a specified intervention. Taxon-removal analyses in existing microbiome models probe the sensitivity of predictions or embeddings [15, 16]. Waypoint evaluates drug-exposure classification and drug degradation prediction [17], but does not demonstrate prediction of post-perturbation community composition. That task requires learning how an introduced organism or compound changes the abundances of community members. Virtual-cell models such as State illustrate how biological representations can be coupled to perturbation information [22]. Extending this principle to microbial communities could help anticipate intended ecological changes and unintended effects of drugs developed against human targets [23, 24]. The challenge is to learn broadly reusable community representations that also supply the biological context needed for intervention-response prediction.

Here we introduce MGM2, a multimodal foundation model pretrained on 1,821,291 MicrobeAtlas samples comprising 225,067 operational taxonomic units (OTUs) clustered at 99% sequence similarity [9]. MGM2 uses DNA language-model representations of OTU sequences as a molecular reference, conditions them on abundance and contextualizes them within each community. Masked abundance modeling learns quantitative relationships among community members, while alignment to sample descriptions links community states to biological settings. The encoder produces both microbial token states and whole-community representations. Task-specific response models combine these frozen features with donor-community or chemical representations to predict intervention outcomes.

We first test whether MGM2 preserves ecological distinctions in later MGnify studies, including projects absent from pretraining. We then examine two forms of intervention: fecal microbiota transplantation (FMT), which introduces another microbial community, and drug exposure, which introduces a chemical compound. A separate recovery task tests what the measured perturbed state reveals about subsequent change. Finally, sparse autoencoders resolve taxonomic and contextual features within microbial token states. MGM2Atlas (https://www.mgm2atlas.com/) provides an interactive view of these features and the observations supporting their interpretations. These evaluations connect the model’s broad representation of microbial communities to its use in predicting and interpreting their responses.

## Results

### Multimodal pretraining builds sequence-grounded community states

An organism’s molecular identity persists across samples, but its ecological context changes. MGM2 represents both by integrating microbial sequence, abundance and sample description (**Fig. 1a**). The pretraining corpus contained 1,821,291 MicrobeAtlas samples and 225,067 OTUs clustered at 99% sequence similarity [9]. Following biological foundation models that initialize entities with pretrained representations [22, 25], MGM2 uses NTv3-650M to encode OTU nucleotide sequences [26]. Projection and abundance conditioning convert these sequence embeddings into sample-specific microbial token inputs. A Transformer then contextualizes the tokens within each community, while semantic alignment relates the whole-community state to sample descriptors. Samples retained at most 768 microbial tokens (**Supplementary Fig. 1a,b**); the median contained 136 non-zero OTUs, with a long tail of larger communities.

**Figure 1.**
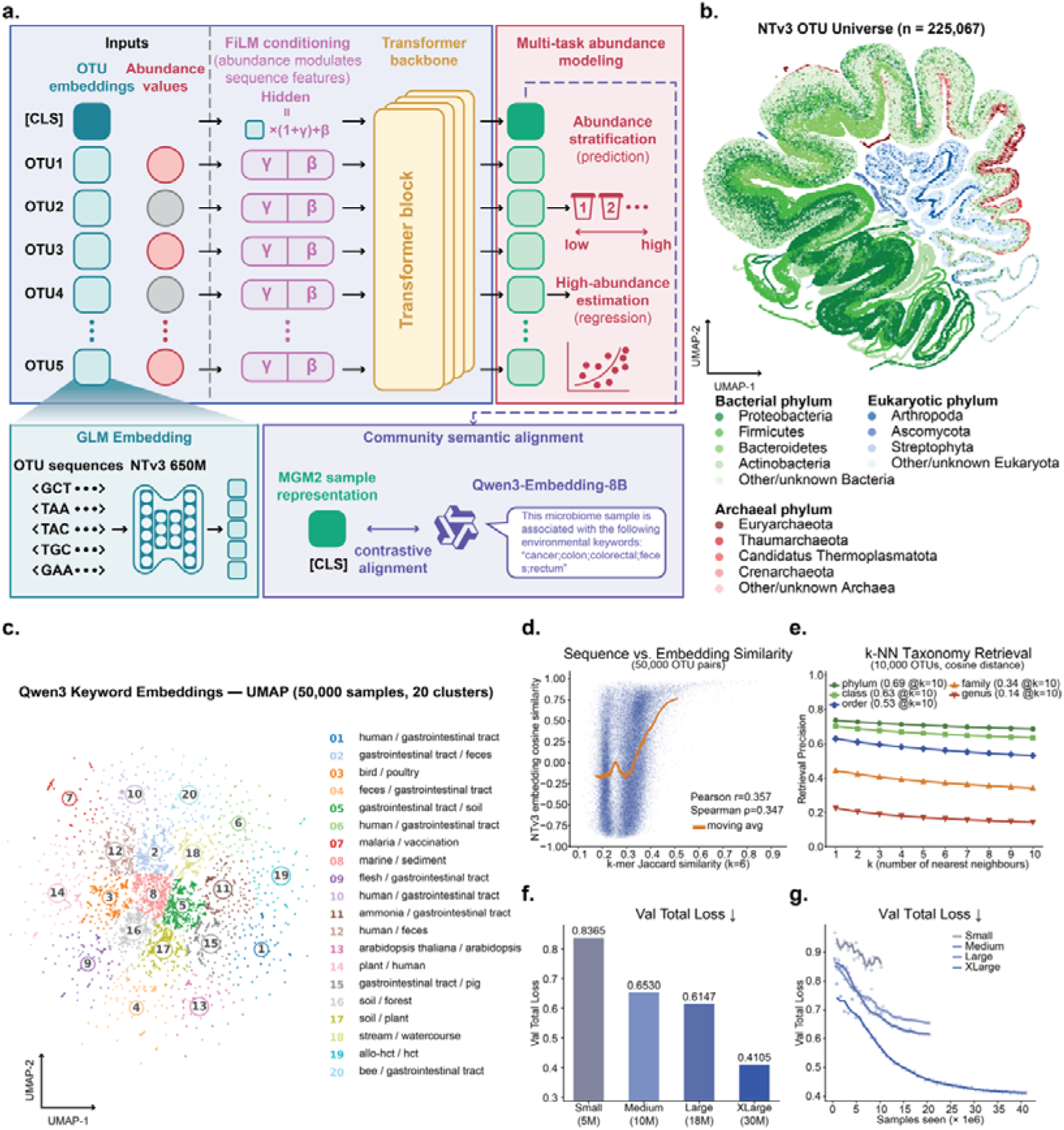
Multimodal architecture and pretraining of MGM2. **a.** NTv3-650M encodes OTU nucleotide sequences. Projection and FiLM abundance conditioning convert the sequence embeddings into microbial token inputs for Transformer contextualization. Gamma and beta denote the feature-wise scaling and shift vectors, respectively. GLM denotes the DNA language model used to encode OTU sequences. Pretraining combines masked abundance reconstruction with optional contrastive alignment to Qwen3-Embedding-8B keyword targets. **b.** UMAP of NTv3 sequence embeddings for the 225,067 OTUs, colored by broad taxonomic group. **c.** UMAP of keyword embeddings for microbiome-associated environmental and disease descriptors. **d.** Relationship between 6-mer Jaccard sequence similarity and NTv3 sequence-embedding cosine similarity across 50,000 OTU pairs. **e.** Taxonomic retrieval precision among nearest neighbors in NTv3 sequence-embedding space. **f.** Best validation total loss for each MGM2 configuration. **g.** Validation loss trajectories during pretraining.

OTU sequence embeddings retained sequence and taxonomic structure before community pretraining. UMAP projections showed broad organization across bacterial, archaeal and eukaryotic phyla (**Fig. 1b**). Across 50,000 randomly sampled OTU pairs, embedding cosine similarity correlated with 6-mer Jaccard sequence similarity (Pearson r = 0.36; Spearman rho = 0.35; **Fig. 1d**). Highly similar sequences had similar embeddings, whereas less similar pairs spanned a broader range of cosine similarities. Taxonomic retrieval precision reached 0.69 at phylum, 0.63 at class and 0.53 at order (**Fig. 1e**). Within-group cosine similarity exceeded between-group similarity from phylum to species, with the largest separation at genus and species (**Supplementary Fig. 1c**). The sequence embeddings thus supplied a taxonomically structured input to community pretraining.

Sample descriptions provide an ecological frame of reference for the molecular and abundance inputs. We contrastively aligned CLS representations with Qwen3-Embedding-8B embeddings of environmental, host and disease-related keywords (**Fig. 1a**). In UMAP space, these embeddings organized into host-associated, aquatic, soil, plant, agricultural, food, livestock and disease-associated contexts (**Fig. 1c**). Language-model embeddings have likewise supplied biological priors for genes and cellular perturbations [27, 28]. In MGM2, sample-level semantic alignment complements token-level abundance reconstruction, linking quantitative community structure to its biological setting.

The joint pretraining objectives improved across the four model configurations (**Table 1**). Validation total loss decreased from 0.84 for Small (5 million parameters) to 0.65, 0.61 and 0.41 for Medium, Large and XLarge (**Fig. 1f**). The latter models contained 10 million, 18 million and 30 million parameters, respectively. Larger configurations converged earlier and had smoother loss trajectories (**Fig. 1g**; **Supplementary Fig. 1e**). From Small to XLarge, abundance-bin accuracy increased from 0.46 to 0.52, masked regression mean squared error fell from 0.27 to 0.23, and regression correlation increased from 0.67 to 0.73. Semantic-alignment loss decreased from 0.60 to 0.21 (**Supplementary Fig. 1d**). Validation grid search selected a shared learning rate, masking probability and semantic-alignment weight (**Supplementary Fig. 1f**). At the selected weight, alignment reduced masked regression error at every scale relative to no alignment, particularly for Large and XLarge (**Supplementary Fig. 1g**). Sample descriptions therefore contributed to learning quantitative community structure as well as supplying a semantic target.

**Table 1.** Architectural configurations of MGM2.

| Model | Hidden size | Layers | Heads | Feed-forward size |
| --- | --- | --- | --- | --- |
| MGM2-small | 224 | 6 | 4 | 896 |
| MGM2-medium | 320 | 6 | 5 | 1,280 |
| MGM2-large | 384 | 8 | 6 | 1,536 |
| MGM2-xlarge | 512 | 10 | 8 | 2,048 |

### Community states preserve ecological distinctions across later studies

Ecological annotation asks whether a newly sampled microbiome can be placed within the diversity described by previous studies [6, 7]. To test this transfer, we evaluated hierarchical biome classification on a study-level temporal MGnify benchmark. Classifiers were trained on studies released before 1 January 2025 and tested on later studies, comprising 2,839 training studies with 291,970 samples and 561 held-out studies with 95,422 samples (**Fig. 2a**). A project-clean subset excluded all test projects represented in MGM2 pretraining, retaining 300 studies and 31,765 samples. This subset tests transfer to projects not encountered during pretraining. Host-associated and environmental communities predominated, alongside a smaller engineered category (**Supplementary Fig. 2a**).

**Figure 2.**
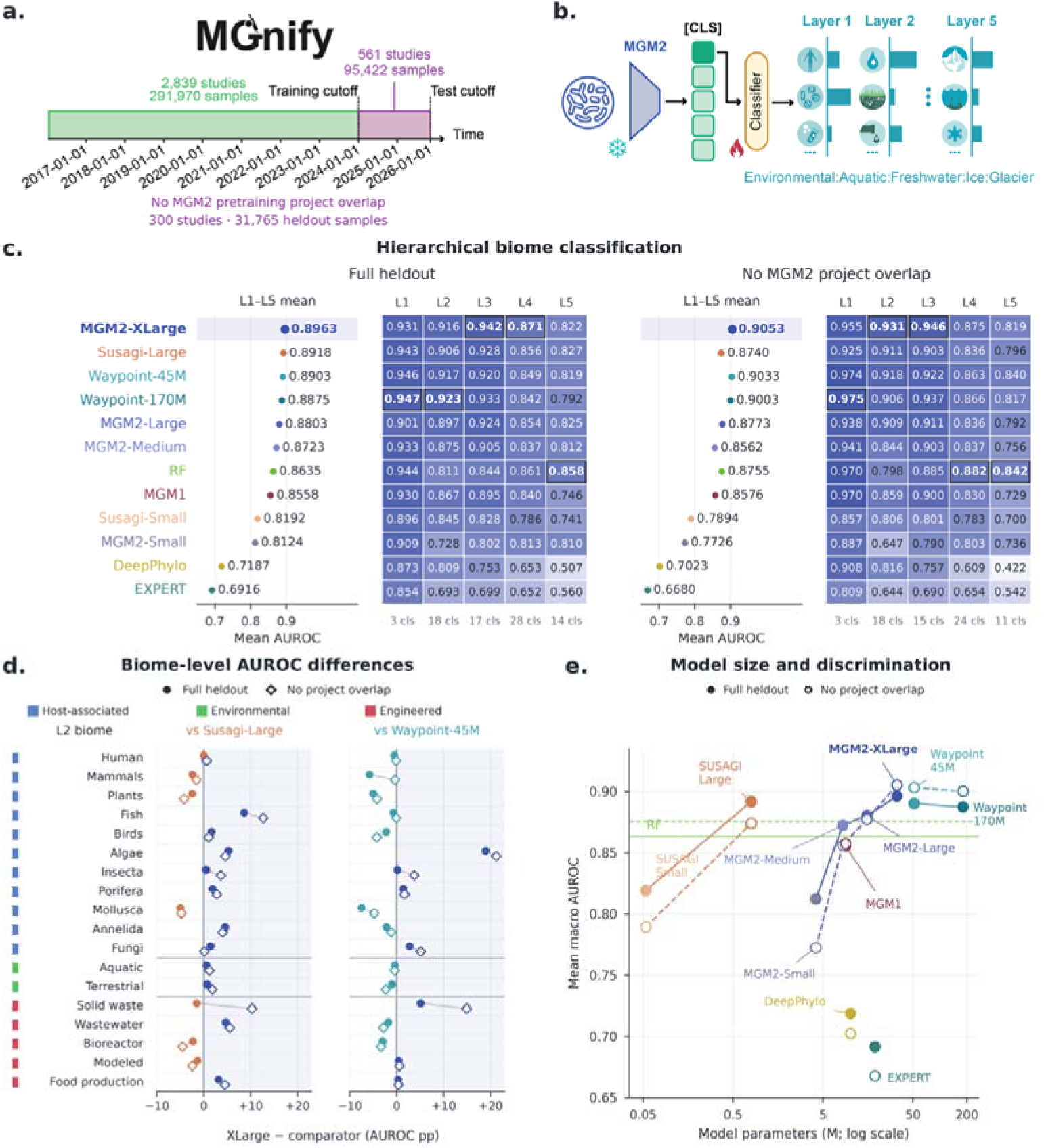
Hierarchical biome discrimination across community-model families. **a.** Study-level temporal split at 1 January 2025: 291,970 training samples from 2,839 studies and 95,422 held-out samples from 561 studies. The project-clean subset contains 31,765 samples from 300 studies after excluding all projects represented in MGM2 pretraining. **b.** Frozen MGM2 CLS representations and level-specific classifiers for the five MGnify biome levels. **c.** Mean macro one-versus-rest AUROC across L1–L5 and level-wise AUROC for 12 models on full and project-clean test sets. Models are ordered by full-set mean AUROC. Each level and population uses identical evaluable samples and shared class definitions across models; class counts appear below columns. Boxed bold cells mark column maxima. Points show descriptive means across levels; effective-run counts are given in **Methods**. **d.** L2 biome-specific AUROC differences between MGM2-XLarge and Susagi-Large or Waypoint-45M, retaining all 18 biomes. Comparators have the highest full-set five-level mean within their displayed families. Differences are in percentage points; positive values favor MGM2-XLarge. Filled circles and open diamonds denote full and project-clean data, respectively; connecting lines identify the same biome. Rows are grouped by parent biome and ordered by full-set support. **e.** Five-level mean AUROC versus model parameter count. Filled circles/solid lines denote full heldout; open circles/dashed lines denote project-clean data. Lines connect configurations within each model family; RF is a horizontal reference. Counts cover specified community models or backbones, with external encoders and classification heads handled as detailed in **Methods**. MGM1 denotes the previous MGM framework.

We held the MGM2 encoder fixed and trained lightweight classifiers on CLS representations at each level of the MGnify hierarchy (**Fig. 2b**). The same pretrained representation was therefore reused from broad biome categories to fine-grained labels. For taxonomic nodes without precomputed sequence-derived embeddings, a taxonomy-guided fallback averaged and normalized embeddings from available descendants (**Supplementary Fig. 2b**). A UMAP comparison with directly precomputed embeddings illustrated the resulting neighborhoods for the 20 most frequent genera (**Supplementary Fig. 2c**).

The frozen community states supported ecological discrimination across the full biome hierarchy. We compared four MGM2 sizes with RF, MGM1, EXPERT, DeepPhylo, Susagi-Small/Large and Waypoint-45M/170M (**Fig. 2c**). MGM2-XLarge had the highest five-level mean macro-AUROC in this comparison: 0.90 on the full held-out set and 0.91 on the project-clean subset. On the full set, its absolute AUROC advantages over Susagi-Large and Waypoint-45M were 0.0045 and 0.0060, respectively. On project-clean data, the corresponding differences were 0.031 and 0.0020. RF reached 0.86 and 0.88, and MGM1 reached 0.86 in both populations. The resulting representation thus retained ecological information across later studies, including projects absent from pretraining.

The hierarchy resolved where these representations were most discriminative (**Fig. 2c**). MGM2-XLarge led the comparison at L3 and L4 on the full held-out set, with macro-AUROCs of 0.94 and 0.87, and at L2 and L3 on project-clean data, with 0.93 and 0.95. Waypoint-170M led at L1 and L2 on the full set and at L1 on project-clean data; RF led at L5 in both populations and at L4 on project-clean data. These complementary rankings place MGM2’s strongest discrimination at intermediate ecological resolutions, where broad habitat categories divide into more specific community settings.

Biome-resolved comparisons reveal ecological variation beneath the hierarchy-wide averages (**Fig. 2d**). Across all 18 L2 biomes, MGM2-XLarge showed gains over Susagi-Large in fish-associated and algal communities and over Waypoint-45M in algal communities; solid-waste communities showed gains over both comparators after project filtering. Other biomes favored the comparator, indicating that the aggregate ranking combines distinct patterns across host-associated, environmental and engineered communities. This resolution links model-level performance to the community types for which ecological annotation is required.

Model size provided a second view of the learned ecological structure (**Fig. 2e**). Five-level mean AUROC increased across the four MGM2 configurations, from 0.81 to 0.90 on the full held-out set and from 0.77 to 0.91 on project-clean data. The parameter-count comparison places this within-family progression alongside Susagi, Waypoint and the other architectures, with RF shown as a reference. Per-class results span the full observed range of label support (**Supplementary Fig. 2e,f**).

The classification results were accompanied by ecological organization within the representation itself. On 1,303 shared project-clean samples, we compared raw abundance with all four MGM2 scales using AvgBIO, which combines clustering agreement with biome labels and normalized cosine-silhouette separation (**Supplementary Fig. 2d**). MGM2-XLarge had the highest AvgBIO at L2–L5, whereas transformed raw abundance scored higher at L1. Thus, MGM2 preserved distinctions at finer ecological resolutions across studies, providing a reusable basis for community analysis.

### Paired community states inform fecal microbiome transplantation outcomes

FMT introduces an entire microbial assemblage into an existing ecosystem. Although clinical use of fecal transfer predates modern microbiome profiling by decades [29], sequence-resolved studies now show how donor and recipient communities contribute to the post-transplant microbiome [30, 31]. This paired-community setting tests whether an encoder pretrained on individual microbiomes can inform intervention outcomes. We used frozen MGM2 representations to predict clinical response and post-FMT composition. Clinical evaluation included 184 labeled events from 178 patients and 17 studies in the primary FMT dataset [30], spanning seven indications with 112 responders and 72 non-responders (**Fig. 3a**). Matched donor, pre-FMT recipient and post-FMT profiles supported the separate composition task.

**Figure 3.**
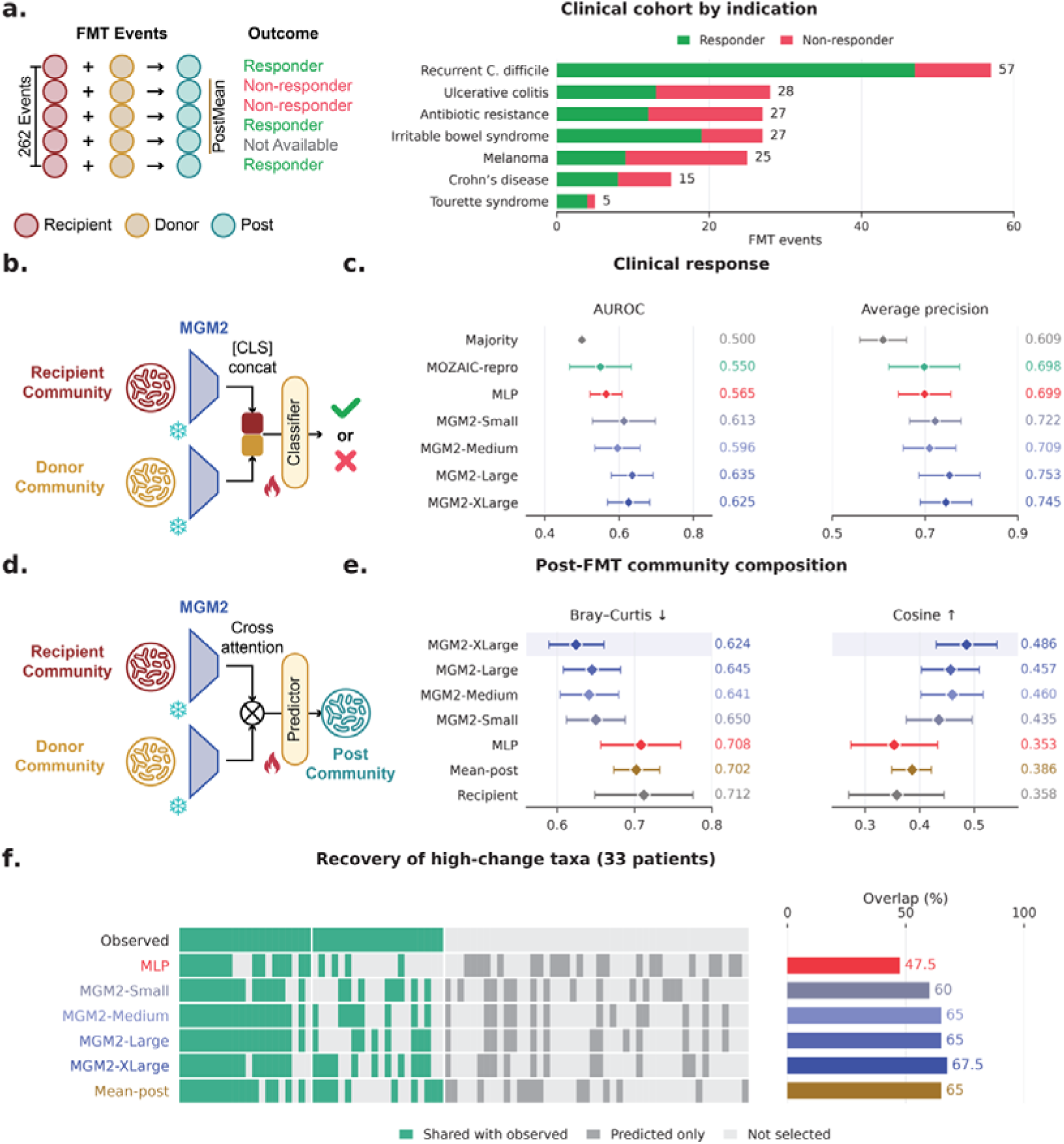
Paired community representations support FMT response and composition prediction. **a.** Clinical cohort of 184 events from 178 patients and 17 studies, including 112 responders and 72 non-responders. Bars show event counts across seven indications. **b.** Response classification from concatenated donor and recipient CLS representations with frozen MGM2 backbones. **c.** Clinical-response AUROC and average precision. Metrics are averaged over three training seeds within each patient-grouped outer fold. Diamonds and error bars show the unweighted mean ± sample SD across five folds. Responder is the positive class. Majority predicts training responder prevalence; MOZAIC-repro uses microbiome inputs only. **d.** Simplified community-prediction architecture. The evaluated head combines cross-attention, timepoint conditioning and gated delta fusion (**Methods**). **e.** Composition prediction for 117 targets from 33 patients, 33 FMT events and 13 studies. The patient shared with training was excluded without retraining. Metrics are averaged over five seeds within target, then over targets within patient and equally across patients. Diamonds show means; error bars show 95% percentile intervals from 2,000 patient bootstrap resamples. Recipient retains pre-FMT composition; Mean-post uses the fixed training-post mean. Lower Bray-Curtis and higher cosine indicate closer agreement. **f.** Recovery of high-change taxa among 755 features after averaging predictions across seeds and profiles within patients. Each set selects the 20 largest positive and 20 smallest negative mean abundance changes, retaining ties and excluding zeros. The union contains 86 displayed taxa. Green denotes observed/shared membership, dark gray predicted-only membership and light gray non-selection. Bars show the fraction of the observed 40-taxon set recovered, without an additional sign-matching requirement.

Paired community representations improved clinical-response discrimination. A lightweight classifier combined donor and recipient CLS states from the same frozen MGM2 backbone (**Fig. 3b**). Models used shared patient-grouped five-fold partitions, four validation candidates per learned model and three training seeds (**Methods**). MGM2-Large achieved the highest mean AUROC (0.64) and average precision (0.75), followed by MGM2-XLarge (0.63 and 0.74; **Fig. 3c**). The abundance MLP reached 0.56 and 0.70, while the microbiome-only MOZAIC reproduction [32], denoted MOZAIC-repro, reached 0.55 and 0.70. All four MGM2 scales exceeded both learned baselines in mean AUROC and average precision.

Predicting post-FMT composition asks how the introduced community reorganizes the recipient microbiome. A time-conditioned head combined recipient and donor token states through cross-attention and gated delta fusion (**Fig. 3d**). We evaluated 117 held-out post-FMT targets from 33 patients, 33 events and 13 studies after excluding the patient shared with training; scores gave equal weight to patients. MGM2-XLarge achieved the lowest mean Bray-Curtis dissimilarity (0.62) and highest mean cosine similarity (0.49) among seven methods, compared with 0.71 and 0.35 for the abundance MLP (**Fig. 3e**). It also improved on retaining the recipient profile or using the mean training-post profile.

The composition predictions recovered taxa undergoing substantial change after transplantation. MGM2-XLarge recovered 27 of the 40 observed high-change taxa (68%), compared with 48% for the MLP (**Fig. 3f**). Large, Medium and Mean-post each recovered 65%, while Small recovered 60%. The sets contained taxa with the largest positive and negative patient-weighted mean abundance changes; overlap measured shared taxon identity. Across 755 taxa, averaged MGM2-XLarge predictions achieved a log_2_ fold-change Spearman correlation of 0.49, compared with 0.45 for the MLP (**Supplementary Fig. 3b**). All MGM2 scales also exceeded the MLP in the three mean per-seed change-recovery metrics (**Supplementary Fig. 3a**).

Model-scale rankings differed across FMT endpoints. Large led clinical-response AUROC and average precision, whereas XLarge led composition agreement and pooled high-change taxon recovery. Large had the highest mean delta cosine (0.65), closely followed by XLarge (0.65). Disease-specific recovery also varied (**Supplementary Fig. 3d**). XLarge, Small and Mean-post each recovered 73% of high-change taxa in recurrent C. difficile, whereas Mean-post exceeded XLarge in antibiotic resistance (53% versus 50%). The scale dependence thus differed between clinical discrimination and the reconstruction of microbial changes.

A separate dataset from Su et al. [32] tested whether paired states could classify the direction of change in donor similarity (**Supplementary Fig. 3c**). Its 511 events formed 375 groups linked by shared recipient samples, which were kept disjoint across training, validation and testing in ten disease-stratified splits. MGM2-XLarge achieved the highest mean AUROC (0.77), compared with 0.66 for MOZAIC-repro, and all four MGM2 scales exceeded the reproduction. Here the positive class denoted non-positive donor-similarity change, a compositional endpoint distinct from clinical response. Across these tasks, the same frozen encoder related donor and recipient communities to both clinical outcomes and microbial reorganization.

### Joint molecular and community representations predict drug responses

Predicting how a compound reshapes a microbiome could help prioritize drugs for desired community changes and identify unwanted ecological effects [23, 24]. This requires a model to connect the exposed community with the chemical structure of the intervention, paralleling the state-and-perturbation formulation of virtual-cell models [22]. We tested this extension of MGM2 using two reprocessed experimental panels from Shi et al. [33]. The drug-perturbation panel contained eight synthetic intestinal communities (SICs), 90 drugs and 720 SIC–drug conditions, represented by 728 treated wells and 16 concurrent DMSO controls (**Fig. 4a**). The controls supplied each SIC’s reference composition. A separate recovery panel followed SIC0 after 23 drugs or DMSO, with two reseeding arms and two replicates, yielding 96 trajectories and 288 measurements across R1–R3. These panels test drug-conditioned composition prediction and recovery from a measured perturbed state.

**Figure 4.**
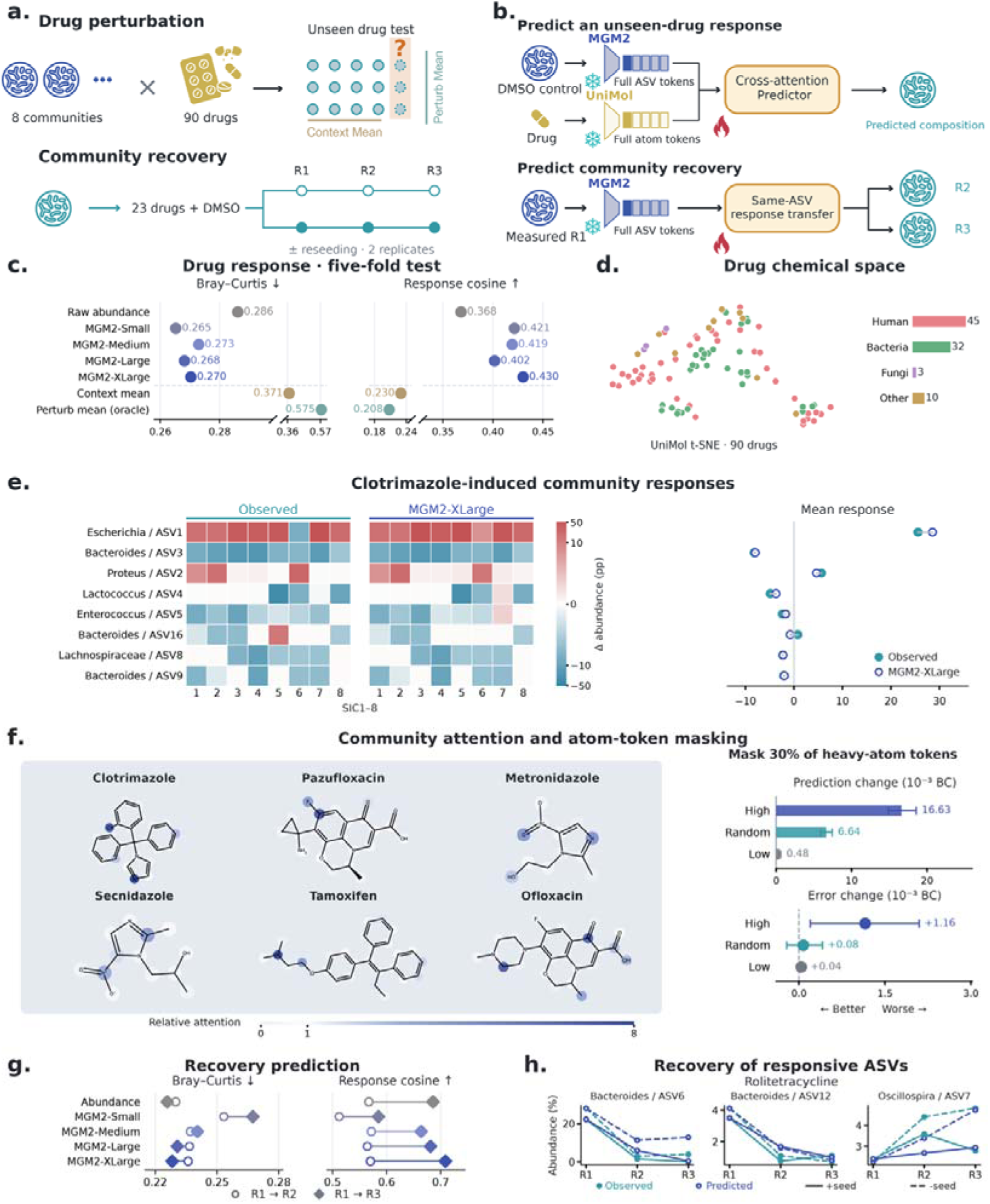
Prediction of drug effects and community recovery. **a.** Perturbation panel comprising eight SICs, 90 drugs, 720 SIC–drug conditions, 728 treated wells and 16 DMSO controls. The separate recovery panel contains 96 trajectories and 288 measurements across R1–R3, from 23 drugs or DMSO with two reseeding arms and two replicates. **b.** Drug prediction combines global MGM2 context and raw abundance with whitened PCA32 ASV queries and frozen Uni-Mol atom representations. Recovery transfers same-ASV responses from measured R1 without drug embeddings. **c.** Outer-test Bray-Curtis dissimilarity and response cosine from nested drug-grouped five-fold evaluation. Wells are scored individually, then averaged within SIC–drug pairs and equally across SICs and drugs. XLarge uses an expanded inner search (145 candidates); other representations use 37 (**Methods**). Context mean uses training drugs within SIC; Perturb mean is an oracle using the held-out drug’s outcomes in other SICs. **d.** Uni-Mol t-SNE and target categories for all 90 drugs. **e.** Observed and predicted Clotrimazole responses relative to DMSO. Clotrimazole has the highest outer-test response cosine; displayed taxa have the eight largest mean absolute observed changes among annotated taxa. Heatmaps use a shared symmetric-log percentage-point scale; the right panel shows SIC-averaged responses. **f.** Community attention for the six drugs with highest response cosine; one denotes uniform heavy-atom attention. At right, prediction shifts and observed-outcome error changes after masking 30% of heavy-atom tokens across all 90 drugs. Random controls average ten equal-sized masks. Whiskers show descriptive 95% drug-bootstrap intervals (2,000 resamples); positive error change indicates worse accuracy. The display fraction was selected after examining 10%, 20% and 30% masking; selection and uncertainty are detailed in **Methods**. **g.** Separate nested five-fold recovery evaluation grouped by initial condition. Open circles and filled diamonds denote direct R1→R2 and R1→R3 predictions. **h.** Selected Rolitetracycline trajectories for three ASVs. Teal/blue denote observed/predicted abundance; solid/dashed lines denote reseeded/unseeded arms. Values average two replicates.

Joint community and molecular representations improved prediction of drug effects on community composition. Frozen MGM2 community and amplicon sequence variant (ASV) states were combined with Uni-Mol drug representations [34] and raw abundance; a global MLP and a gated ASV-to-atom attention branch predicted compositional changes relative to DMSO (**Fig. 4b**). Nested five-fold evaluation held out all observations of each test drug from model fitting and inner selection, covering all 90 drugs in the same eight communities (**Methods**). MGM2 models achieved Bray-Curtis dissimilarities of 0.27 and response cosines of 0.40–0.43, compared with 0.29 and 0.37 for the raw-abundance attention comparator (**Fig. 4c**). Small had the lowest dissimilarity, whereas XLarge had the highest response cosine. Fold-level scores show the variation across held-out drug groups (**Supplementary Fig. 4a**).

Conditioning on both inputs outperformed predictions based on average responses. Context mean, which averaged training-drug outcomes within the same SIC, gave Bray-Curtis dissimilarity 0.37 and response cosine 0.23. Perturb mean, an oracle reference using the held-out drug’s measured outcomes in the seven other SICs, gave 0.58 and 0.21 (**Fig. 4c**). The drug collection included compounds targeting human, bacterial and fungal processes (**Fig. 4d**), extending the task beyond antibiotic exposure. In the selected Clotrimazole example, MGM2-XLarge captured increased Escherichia/ASV1 and decreased Bacteroides/ASV3 across SICs, as well as community-dependent changes in other taxa (**Fig. 4e**). This illustrates how chemical conditioning and community state jointly determine a predicted response.

The molecular interface also allowed us to examine which chemical inputs influenced the predictions. For XLarge, an expanded capacity and regularization search selected the attention-branch configuration within each outer fold; **Supplementary Fig. 4b** shows all tested profiles and the inner-selected pipeline. Attention rankings were largely shared across microbial queries: 96% of the weighted ASV queries had a leave-one-out rank correlation of at least 0.8 with the other ASVs in the same SIC and drug (**Supplementary Fig. 4c**). This agreement supported a community-level summary, obtained by abundance-weighting ASV attention within SICs and then averaging across SICs.

Masking highly attended molecular tokens produced larger changes in the fitted predictor than masking random or low-attention tokens. Across all 90 held-out drugs, excluding the highest-attention 30% of heavy-atom tokens shifted predictions by 0.017 Bray-Curtis units on average, compared with 0.0066 for random masks and 0.00048 for low-attention masks (**Fig. 4f**). The high-minus-random increase in observed-outcome error was 0.0011 (descriptive 95% drug-bootstrap interval, 0.00029–0.0019). Effects varied across drugs, with positive differences for 52 and negative differences for 38 (**Supplementary Fig. 4d**). These interventions locate model sensitivity at the community–molecule interface. The six attention maps illustrate drugs with high response cosine; selection of the displayed masking fraction and the full intervention procedure are described in **Methods**.

We next asked whether the perturbed community state alone could inform subsequent recovery. Longitudinal studies have shown that recovery after antibiotic exposure can be incomplete and individualized [35]. Here, a same-ASV response-transfer model used measured R1 states to predict R2 and R3 separately within each reseeding arm, without drug embeddings (**Fig. 4b,g**). Nested five-fold evaluation grouped all arms and replicates of each initial condition together. At R3, MGM2-XLarge had the highest response cosine (0.71 versus 0.69 for abundance), while Medium led at R2. The abundance comparator had the lowest Bray-Curtis dissimilarity at both horizons. Selected Rolitetracycline trajectories showed recovery of Bacteroides/ASV6, Bacteroides/ASV12 and Oscillospira/ASV7 in both reseeding arms (**Fig. 4h**). MGM2 thus informed the direction of later recovery, complementing its use in predicting drug-induced reorganization.

### Sparse features reveal how microbial identity and ecological context coexist

A transferable community representation should capture both who an organism is and the context in which it occurs. To examine how MGM2 organizes this information, we decomposed microbial token states with sparse autoencoders (SAEs), extending an approach used in language models [36] and biological models including ESM and Evo 2 [12, 37, 38]. A BatchTopK SAE [39] was trained on OTU-token states from 500,000 microbiome samples (**Fig. 5a**). The selected dictionary contained 4,096 features, with an average of 64 active features per token across a batch. It combined high reconstruction performance with broad feature usage: 4,094 features had positive activations in the train/validation interpretation set (**Supplementary Fig. 5a**).

**Figure 5.**
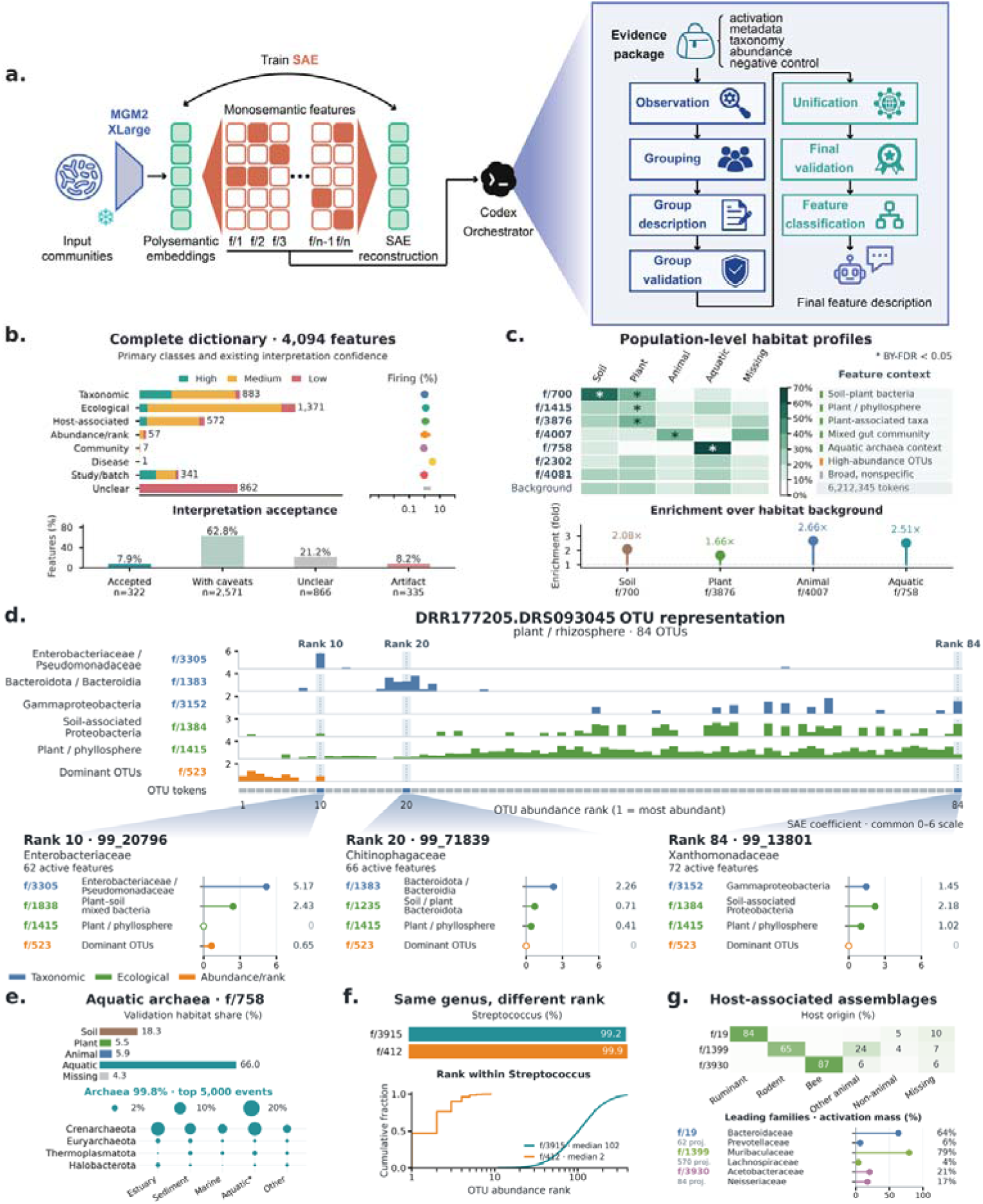
Biological and contextual structure in the sparse feature dictionary. **a.** BatchTopK SAE decomposition and evidence-based feature interpretation. **b.** Primary classes, confidence and acceptance for all 4,094 interpreted non-dead features. Firing dots and intervals show median and interquartile range across features on 6,212,345 validation OTU-token events. Class and acceptance are separate fields. **c.** Habitat shares among each feature’s positive validation events, including missing metadata, with the all-token background below. Colors share a 0–70% scale. Stars mark positive enrichment with study-cluster Benjamini–Yekutieli adjusted P < 0.05 (19,999 draws, 8,959 studies, 16,376 hypotheses). Lower points show token-weighted enrichment over habitat background. Contrasts 2302 and 4081 were curated after testing. **d.** Complete 84-token plant/rhizosphere sample, ordered by abundance rank. Six tracks have independent raw scales; selected expansions at ranks 10, 20 and 84 share a 0–6 coefficient scale and retain zeros. **e.** Feature 758: full-validation habitat shares above; taxonomic event counts and phylum-by-habitat activation mass in a separate retained top-5,000 event set below. Bubble denominators include the entire selected set. **f.** Streptococcus fractions in separate top-5,000 sets for features 3915 and 412; lower curves show ranks among their Streptococcus events. **g.** Host-origin event shares and leading named-family activation-mass fractions in separate 5,000-event sets. All mass denominators retain unassigned taxa; project counts are 62, 570 and 84. Local examples use selected coordinates and retained high-activation events; population habitat estimates are shown in c.

The sparse dictionary resolved several levels of biological description (**Fig. 5a,b**). Deterministic evidence packages linked all 4,094 active features to taxonomy, abundance, metadata and matched controls; Codex assessed candidate descriptions against these records. Primary classes included ecological niche (1,371 features), taxonomic identity (883), host-associated context (572), abundance/rank (57), community interaction or co-occurrence (7), disease-associated context (1), study-associated variation (341) and unclear interpretation (862). A separate confidence assessment accepted 322 descriptions, accepted 2,571 with caveats, retained 866 as unclear and classified 335 as artifact-associated. MGM2Atlas (https://www.mgm2atlas.com/) makes the feature descriptions, evidence and confidence assessments available for interactive exploration.

Population-level habitat profiles connected selected features to their ecological distributions across 6,212,345 validation OTU-token events from 8,959 studies (**Fig. 5c**). Habitat shares were calculated over all positive activations of each feature, retaining missing metadata. Feature 758 had 66% aquatic activations, compared with 26% aquatic tokens in the background, a 2.5-fold enrichment. Plant-associated features 1415 and 3876 had plant shares of 25% and 34%, respectively, compared with 20% in the background. Study-cluster tests corrected across 4,094 features and four habitats identified six enriched pairs among those displayed, including both plant features. Broad feature 4081 closely followed the habitat background. These profiles distinguish habitat enrichment from widespread activation; **Supplementary Fig. 5b,c** shows within-habitat firing distributions and the validation population composition.

Individual microbial tokens combined taxonomic, ecological and abundance-related features within the same community (**Fig. 5d**). In a complete plant/rhizosphere sample containing 84 OTUs, taxonomic coordinates varied across abundance-ranked tokens, while feature 1415 was active in 75 tokens and dominant-OTU feature 523 was active at ranks 1–7 and 10. Expansions at ranks 10, 20 and 84 showed these coefficients together, including zero values. The sparse representation therefore expressed microbial identity alongside position and shared context within a community.

Selected feature views showed how this separation operates within and across taxa. In the retained top-5,000 events for aquatic-associated feature 758, 4,991 were archaeal and spanned several phyla (**Fig. 5e**). Features 3915 and 412 both concentrated on Streptococcus, but their Streptococcus events had median abundance ranks of 102 and 2, respectively (**Fig. 5f**). A shared genus identity thus occupied different sparse coordinates according to abundance context. Host-associated features 19, 1399 and 3930 connected ruminant, rodent and bee contexts to different taxonomic families across 62, 570 and 84 projects (**Fig. 5g**). These event-level views identify the organisms and compositional settings contributing to the population patterns.

Sparse features also exposed study-associated structure. Of feature 973’s 5,000 retained events, 4,999 came from ERP022452, and all were WGS dog samples (**Supplementary Fig. 5d**). Nearly all activation mass was concentrated in that study, compared with top-study shares of 18% for bee-associated feature 3930 and 5.8% for rodent-associated feature 1399. In a complete 241-token sample from ERP022452, feature 973 was active in 235 tokens alongside taxonomic, gut-context and rank-related coordinates (**Supplementary Fig. 5e**). The feature atlas therefore reveals both biological associations and their concentration in particular sources, making the basis of an interpretation accessible for further testing.

## Discussion

MGM2 brings molecular identity and ecological context into a shared representation of microbial communities. Sequence and microbiome foundation models have established the value of pretraining at these respective levels [11–17]. MGM2 connects them through DNA language-model representations, quantitative abundance conditioning and alignment to biological descriptions. The descriptions improved masked abundance reconstruction, showing that semantic context contributed to learning quantitative community structure. Transfer to later MGnify projects absent from pretraining demonstrated that the resulting frozen states retained ecological distinctions across studies. This multimodal representation supplies a common starting point for community analysis and response models with different intervention inputs.

FMT shows how a foundation model trained on individual communities can support prediction when two communities meet. Strain-tracking studies established the importance of donor and recipient context for engraftment [30, 31], and targeted models now use paired microbiomes to inform matching [32]. MGM2 carries knowledge learned across diverse microbiomes into this setting through a shared frozen encoder. Paired representations supported clinical-response classification, post-transplant composition prediction and recovery of high-change taxa. Their different model-scale rankings indicate that clinical response and microbial reorganization draw on related but distinct information. Learning a common representation allows these endpoints to be studied together while retaining their biological meaning.

Drug perturbation extends this approach from an introduced community to an intervention represented by molecular structure. Community interactions can protect or sensitize microorganisms to drugs [24, 33], making the exposed community an important part of the prediction. Coupling MGM2 states to Uni-Mol representations [34] supported prediction of drug-induced compositional changes in the evaluated SICs, while the recovery task used measured perturbed states to predict subsequent change without chemical input. This goes beyond assigning drug-related properties to a microbiome: the output is a predicted community composition conditioned on an intervention or an earlier state. The connection to virtual-cell modeling [22] is therefore operational, linking a biological state, an input and a predicted response. Broader compound and community coverage could make this approach useful for microbiome-aware drug discovery, including screening for desired compositional changes and ecological side effects.

The sparse feature dictionary makes the model’s representation of context inspectable. Taxonomic, habitat-associated and abundance-related features co-occurred within individual microbial token states; the Streptococcus features showed how one genus could occupy different coordinates according to abundance rank. Population profiles, matched controls and study-source distributions anchor these descriptions to observable data, and MGM2Atlas provides access to that evidence. Attention interventions in the drug predictor examine a complementary level: which molecular inputs influence a response prediction. These analyses turn distributed representations into specific hypotheses about community organization and chemical sensitivity that can be tested experimentally.

MGM2’s current limitations center on functional resolution and transfer across intervention settings. Connecting sequence-derived community states to metabolic activity requires strain-resolved functional measurements. Prospective FMT cohorts can establish how paired representations inform donor selection, while drug-response evaluation in additional communities and experimental systems can test transfer beyond the in vitro communities studied here. For recovery, improved response direction remains distinct from lower absolute composition error. Controlled ecological and chemical interventions can further determine which sparse associations and model-sensitive molecular inputs reflect biological mechanisms.

MGM2 establishes a multimodal foundation for connecting microbial community structure with responses to intervention. Knowledge learned from large-scale observational profiles transfers to ecological characterization and, when coupled to the relevant community or chemical inputs, to response prediction. The sparse feature dictionary makes the taxonomic and ecological content of this shared representation accessible to inspection. These capabilities bring the description of a microbiome and the prediction of its reorganization into a common modeling framework, providing a basis for investigating how starting community states shape the outcomes of intervention.

## Methods

### Construction of sequence-aware microbiome samples

MGM2 pretraining used 1,821,291 MicrobeAtlas profiles comprising 225,067 OTUs clustered at 99% sequence similarity [9]. We converted raw counts to within-sample relative abundances and removed zero-abundance features. Each retained OTU was linked to a fixed NTv3-650M embedding of its nucleotide sequence. A CLS token provided a community-level representation. Samples retained at most the 768 most abundant microbial features. The stored dataset contained relative abundances; log transformation and masking were applied during training.

### Abundance preprocessing and binning

Relative abundances were converted to a bounded log-abundance channel. Zeros remained 0. Positive values were floored at 1 × 10^−4^, transformed as log_10_(x) + 2 and clipped to [−2, 2]. Sample-specific quintiles defined five ordered positive-abundance labels. Continuous regression supervision was restricted to masked tokens in the two highest abundance states, alongside discrete-bin supervision of masked positions.

### MGM2 architecture

MGM2 uses a pre-layer-normalized Transformer encoder to contextualize microbial tokens within a community [40]. A projection maps the 1,536-dimensional NTv3-650M embeddings of OTU nucleotide sequences into the MGM2 hidden space. Feature-wise linear modulation (FiLM) incorporates sample-specific abundance before contextualization [41]. The abundance conditioner produces gamma and beta vectors that scale and shift each projected sequence embedding to form an initial token state. The learned CLS embedding is not abundance-conditioned. Contextualized microbial tokens support abundance reconstruction, while the CLS state is used for semantic alignment and community-level prediction.

Let *e_i_* denote the fixed NTv3 embedding of OTU sequence *i* and *a_i_* the corresponding transformed abundance. The abundance conditioner produced *γ*(*a_i_*) and *β*(*a_i_*), and the initial token state was

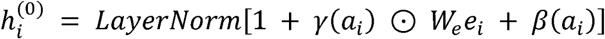

followed by dropout and Transformer contextualization. No positional embeddings were added. The encoder was therefore permutation equivariant over microbial tokens, while the CLS state provided a permutation-invariant community summary apart from padding masks.

### Community-semantic targets

Curated environmental, host and disease-related descriptors were encoded with Qwen3-Embedding-8B to generate 4,096-dimensional sample-level targets. Targets were matched to microbiome profiles using normalized sample identifiers. Samples without matched targets contributed to masked abundance pretraining but not to semantic alignment.

### Pretraining objectives

Pretraining combined masked abundance modeling with optional community-semantic alignment. We selected 15% of non-CLS microbial positions for abundance supervision while retaining their microbial identities. At these positions, an 80/10/10 corruption scheme replaced abundance with a mask value, substituted a random within-sample abundance or retained the original value. The model recovered discrete abundance bins at masked positions and continuous log-abundances for tokens in the two highest abundance states. Bin prediction used focal loss, and continuous prediction used mean squared error. For samples with matched keyword targets, a symmetric in-batch contrastive loss aligned the CLS representation with the corresponding Qwen3-Embedding-8B embedding [42].

Masked abundances were represented by −3, outside the observed transformed range. Let *M* denote masked microbial positions and *M*_ℎ_ the subset assigned to the two highest abundance states. For each *i* in *M*, *p_i_* was the predicted probability of the true abundance bin, *a_i_* was the transformed abundance target and *â_i_* was its prediction.

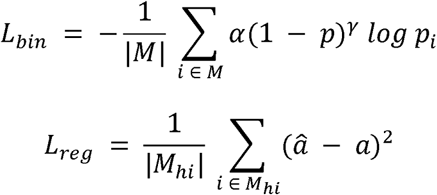

For semantic alignment, B denoted the n samples in a mini-batch with matched Qwen3 targets. A learned linear projection mapped each CLS state to the 4,096-dimensional target space. The projected CLS vector and Qwen3 target were then L2-normalized to *z_i_* and *q_i_*, respectively, and pairwise logits were defined by

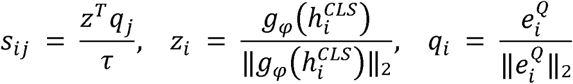

The matching sample formed the positive pair and all other valid samples in the mini-batch formed negatives. Writing S = [*s_i_*_,*j*_] for the similarity matrix, symmetric alignment was computed in both directions:

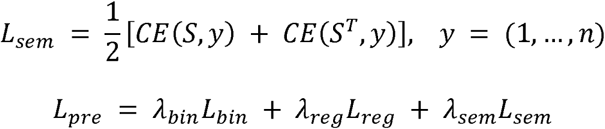

We used α = 0.25 and γ = 2 for focal loss, temperature τ = 0.07, and objective weights of 1.0, 0.3 and 0.1 for bin classification, regression and semantic alignment, respectively. Loss terms without valid supervised positions were set to zero; in particular, semantic alignment required at least two matched samples in a mini-batch.

### Training and scaling configurations

Scaling experiments used a shared recipe selected by validation grid search. Models were optimized with AdamW at a peak learning rate of 5 × 10-5 and weight decay 0.01. The learning rate increased over 1,000 warmup steps and then followed cosine decay to 10% of its peak value. Gradients were clipped at 1.0, training used bfloat16 precision, the random seed was 42 and the data were divided into 90% training and 10% validation samples. Training continued for at most 50 epochs, with early stopping after five validation epochs without improvement and checkpoint selection by minimum validation total loss. Architectural configurations are listed in **Table 1**. Batch sizes were 1,024, 512, 512 and 256 for Small, Medium, Large and XLarge, respectively. All scales used masking probability 0.15 and loss weights of 1.0, 0.3 and 0.1 for abundance-bin classification, continuous abundance regression and Qwen3 alignment, respectively.

### Sequence and semantic embedding analyses

We evaluated NTv3 embeddings of OTU nucleotide sequences independently of MGM2 pretraining. UMAP [43] visualized these sequence embeddings for the 225,067 OTUs. For 50,000 randomly sampled OTU pairs, we compared 6-mer Jaccard sequence similarity with embedding cosine similarity using Pearson and Spearman correlations. Taxonomic retrieval used k = 10 neighbors at ranks from phylum to species. Precision was the fraction of neighbors sharing the query label. We also compared within-label and between-label cosine-similarity distributions at each rank. Qwen3 keyword embeddings were visualized by UMAP to examine their organization by host and environmental descriptors.

### Study-level temporal MGnify benchmark

MGnify data were downloaded on 13 April 2026. For downstream biome classification, MGnify studies released before 1 January 2025 were assigned to training, and studies released after the cutoff were assigned to testing. The model-input training set contained 2,839 studies with 291,970 samples; the full held-out set contained 561 studies with 95,422 samples. Studies were defined by unique ERP, SRP or DRP project accessions, with all downstream sample identifiers mapped to projects.

The downstream training and test study sets were disjoint. A project-clean test subset excluded all projects overlapping the MGM2 pretraining corpus, retaining 300 studies and 31,765 samples. Cohort counts precede level-specific filtering for evaluable labels (**Fig. 2a,c**).

MGM2 parameters were frozen, and level-specific lightweight classifiers were trained on CLS community representations. Classification performance was summarized by macro one-versus-rest AUROC at each of five biome levels (L1–L5). Within each level and test population, all models used identical evaluable sample identifiers and shared classes. Unknown labels and classes without common probability outputs were excluded. Five-level mean AUROC was the equal-weight arithmetic mean of the five level-wise values. Per-class AUROC was also examined against held-out class support (**Supplementary Fig. 2e,f**).

The comparison comprised RF, EXPERT [44], DeepPhylo [45], the previous MGM framework [16] (MGM1 in the figures), Susagi-Small/Large [18] and Waypoint-45M/170M [17], alongside four MGM2 sizes. RF used sparse abundance features aligned by OTU identifier strings and 100 trees. Susagi used frozen DNA-only community checkpoints with genus-aggregated ProkBERT representations; Waypoint used its genus-token representation pipeline. Both external model families used MLP classification heads trained with cross-entropy loss and AdamW (learning rate 10^−4^, weight decay 0.01, batch size 2,048), with validation-loss early stopping (patience 5; maximum 100 epochs).

RF, MGM1, EXPERT, DeepPhylo and the displayed Susagi and Waypoint checkpoints each had five effective prediction runs per hierarchy level; the external-model seeds were 11, 22, 33, 44 and 55. MGM2 had one effective prediction run per configuration and level, except Medium at L4, which had two. Duplicate prediction outputs were collapsed before averaging. Reported AUROCs are means of run-level metrics, not metrics from an ensemble of probabilities. The five-level summary gives equal weight to L1–L5. The L2 comparison includes all 18 evaluable biomes and uses Susagi-Large and Waypoint-45M, the checkpoints with the highest full-set five-level mean within the two external families shown in **Figure 2**. Differences are 100 × (MGM2-XLarge AUROC − comparator AUROC), in percentage points. Biomes are grouped by L1 parent and ordered by full-set sample support within each group.

Parameter counts refer to the community model or specified backbone. MGM2 counts exclude the Qwen3 projection; MGM1 and DeepPhylo use the mean count across their five level-specific models, including the frozen phylogenetic embedding for DeepPhylo; EXPERT uses its saved-model variables. Susagi counts the DNA-only community model (53,641 and 798,801 parameters), excluding the separate ProkBERT encoder and downstream MLP. Waypoint counts unique backbone parameters, including embeddings and the tied output head once (51,766,272 and 181,558,272 parameters for the checkpoints named 45M and 170M), excluding the downstream MLP. **Figure 2e** therefore compares these specified model components rather than end-to-end computational cost. Lines connect the observed configurations within each family.

We evaluated representation-space organization separately from classification using cached frozen embeddings shared by raw abundance and all four MGM2 scales. This project-clean subset contained 1,303 samples. AvgBIO was defined as

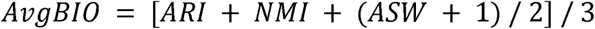

where ARI denotes the adjusted Rand index, NMI denotes normalized mutual information with arithmetic normalization, and ASW denotes the mean cosine silhouette. Features were L2-normalized, and K-means used 20 initializations with random seed 11. The raw-abundance baseline used log1p-transformed abundances followed by a 50-component singular value decomposition refitted on the same subset. Sample selection was balanced by L1 before project filtering, which changed the class proportions in the final subset.

For taxonomic nodes without precomputed sequence-derived embeddings, available descendant embeddings were averaged and L2-normalized (**Supplementary Fig. 2b**). A UMAP diagnostic compared fallback and directly precomputed embeddings for the 20 most frequent genera (**Supplementary Fig. 2c**).

### FMT datasets and prediction tasks

The primary FMT dataset contained 22 studies, 295 annotated FMT-related events, 1,486 samples and 755 microbial features after preprocessing. Metadata distinguished actual FMT from placebo events. Matched donor, pre-FMT recipient and post-FMT profiles yielded 228 FMT events with 700 post-FMT targets for community prediction. Clinical response evaluation used 184 labeled FMT events from 178 study-qualified patients and 17 studies, comprising 112 responders and 72 non-responders. Clinical analyses used shared patient-grouped five-fold partitions rather than a single fixed test split. Community prediction retained its fixed split of 581 training and 119 held-out targets. Excluding two timepoints from the patient shared with training left 117 targets from 33 patients for **Figures 3e,f** and **Supplementary Fig. 3a,b,d**.

### MGM2 clinical response prediction

Clinical response prediction used donor and recipient profiles collected before FMT. A shared frozen MGM2 backbone encoded each community, and a classifier mapped concatenated CLS states through a hidden layer, GELU, dropout and two-class output. Softmax yielded responder probabilities. Baselines were an abundance MLP, microbiome-only MOZAIC-repro and the training-prevalence Majority predictor. All models used the same five indication-stratified outer folds and inner validation partitions, grouped by study-qualified patient identifier. Patients, events and recipient samples were disjoint across partitions; studies and donors could be shared.

Each learned model received four candidate configurations per fold at seed 0. Selection used validation AUROC, followed by validation log loss and configuration name to resolve ties. Selected configurations were retained for seeds 11 and 22. Candidate counts were matched, with model-specific optimization and checkpoint rules as described below. This comparison followed an earlier evaluation of the same cohort. AUROC and average precision were calculated per fold and seed, averaged across seeds within each fold, then summarized by the unweighted five-fold mean and sample SD. Pooled out-of-fold and threshold-dependent metrics were summarized separately.

Outer folds used study-qualified patient identifiers from the combined response/community metadata. Sorted identifiers were shuffled within indication and assigned cyclically using seed 20260909. Validation selected assignment zero from a ten-way indication-stratified patient assignment within the remaining training pool, using seed 20260909 + 1000 + fold index. Saved event memberships defined the exact partitions. Neither response labels nor model outcomes were used to optimize these splits.

The four MGM2 candidates used learning rate 10^−4^ with frozen-backbone training-mode dropout, or 3 × 10^−5^, 10^−4^ and 3 × 10^−4^ with the backbone in evaluation mode. MLP candidates used rates 3 × 10^−4^, 10^−3^ and 3 × 10^−3^ with dropout 0.2, plus 10^−3^ with dropout 0.1. MOZAIC-repro used rates 10^−4^, 3 × 10^−4^ and 10^−3^ with dropout 0.2, plus 3 × 10^−4^ with dropout 0.1. Selected seed-0 checkpoints reproduced saved validation probabilities before test inference. Average precision used average_precision_score rather than trapezoidal precision-recall area. Accuracy, balanced accuracy and macro F1 used a 0.5 threshold.

For response prediction,

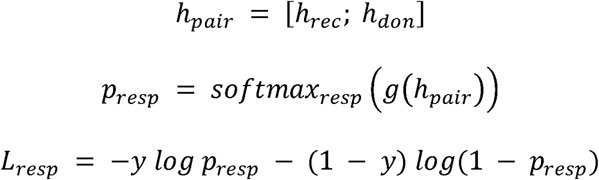

where ℎ*_rec_* and ℎ*_don_* are the recipient and donor CLS embeddings, *g* is the classifier and y denotes responder status.

The clinical MGM2 head mapped concatenated donor and recipient CLS states of dimension 2H through a linear H-dimensional layer, GELU, dropout and a two-class output. Stage-2 backbone weights were frozen. Training used cross-entropy, AdamW, weight decay 0.01, batch size 16 and at most 80 epochs. Checkpoints minimized validation loss with early-stopping patience 12; learning rates and backbone modes followed the candidates above.

The clinical MLP concatenated transformed recipient abundance, donor abundance and donor-minus-recipient features, standardized using training data only. Two 256-unit hidden layers used LayerNorm, GELU and dropout before two-class output. Training used class-balanced cross-entropy, AdamW, weight decay 10^−4^, batch size 16 and at most 300 epochs, with minimum validation-loss selection and patience 30. MOZAIC-repro used microbiome-only DenseNet1D with five layers, initial filters and growth rate 32, and kernel size 5. Donor/recipient standardization was fitted on training data only. Training used binary cross-entropy with explicit L2 penalty 10^−6^, Adam (β₁ = 0.9, β_2_ = 0.999, ε = 10^−8^), batch size 16 and 30 epochs. The maximum validation-AUROC checkpoint was retained.

### MGM2 post-FMT community prediction

Post-FMT community prediction used frozen MGM2 encoders for recipient and donor profiles. Sequence-derived embeddings and transformed abundances formed the microbial token inputs. Recipient states queried donor states and a projected post-FMT timepoint token through cross-attention. Delta fusion combined recipient states, attention output, donor-minus-recipient hidden-state differences and projected abundance differences. A sigmoid gate modulated the update, which was added residually to recipient states and normalized before a token-wise abundance-delta head. Predicted deltas were added to recipient abundances. The objective included errors in transformed post-FMT abundance and transformed abundance change, with weight 0.5 for the latter.

Let *x_rec_* and *x_post_* denote transformed pre-FMT recipient and post-FMT abundance profiles. The donor-conditioned head estimated Δ*_θ_*(*H_rec_*, *H_don_*, *t*), giving:

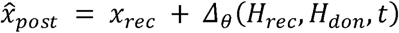

Training minimized:

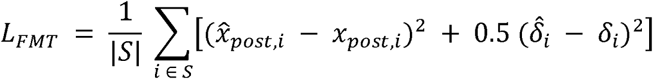

over the supervision mask S. Here, δ and its predicted counterpart denote transformed abundance changes relative to the recipient profile. Prediction export restored relative abundances, applied a raw-zero threshold of 5 × 10^−5^ and renormalized each profile.

Re-evaluation used these saved raw predictions, aligned by target and feature identifiers, without repeating the transformed-output conversion.

Community prediction reused saved MGM2 delta-fusion and MLP delta-loss runs with seeds 11, 22, 33, 44 and 55. MGM2 used frozen backbones, AdamW, learning rate 10^−4^, weight decay 0.01, batch size 32 and at most 100 epochs. A 10% validation split was grouped by FMT event identifier; checkpoints minimized validation loss. The community MLP used three 512-unit hidden layers, dropout 0.15, batch size 64, learning rate 10^−3^ and weight decay 10^−4^. It trained for at most 300 epochs with patience 30 and event-grouped validation. Both objectives combined transformed-post abundance MSE with transformed-delta MSE weighted by 0.5. These runs were separate from the equal-candidate clinical comparison.

### FMT baselines and evaluation metrics

Community baselines comprised Recipient, which retained pre-FMT composition; Mean-post, the fixed mean training-post profile; and an abundance MLP trained with post-abundance and delta losses. Bray-Curtis dissimilarity and cosine similarity were calculated from normalized, feature-aligned profiles for each target and seed. Metrics were averaged across seeds within target, across targets within patient and then equally across patients. Percentile 95% confidence intervals used 2,000 patient bootstrap resamples with seed 20260909. The patient-exclusion subset was evaluated from saved predictions without retraining.

Change recovery was assessed relative to pre-FMT recipient profiles across all 755 taxa. For each seed, patient-weighted mean profiles defined taxon log_2_ fold changes with pseudocount 10^−6^. Delta cosine was computed within targets and then aggregated by patient. Top-Δ overlap measured recovery of the observed 20 largest positive and 20 smallest negative mean changes, retaining ties and excluding zeros. **Figure S3a** reports mean ± sample SD across seeds. For **Figure S3b**, predictions were averaged across seeds before forming patient-weighted profiles and calculating descriptive Spearman correlations.

**Figures 3f and S3d** used averaged predictions to select high-change taxa in the complete cohort and within all seven indications, respectively. Overlap was the observed/predicted intersection size divided by observed set size, without an additional direction-matching criterion. Patients received equal weight within each population, and Mean-post remained fixed from training. All observed sets contained 40 taxa; predicted sets retained boundary ties. These descriptive sets were not selected by differential-abundance significance tests.

### Supplementary donor-similarity evaluation

The supplementary donor-similarity task used 511 events from the Su et al. dataset. Events linked by shared recipient pre- or post-FMT samples formed 375 connected groups, including longitudinal sample reuse across roles. These were recipient-linked groups rather than verified patient identities. Ten disease-signature-stratified splits, using seeds 20260910–20260919, assigned 270/30/75 groups to training/validation/test. Test sets contained 91–110 events, with recipient samples disjoint across partitions. The positive class, similarity_decreased, indicated a non-positive change in donor similarity from pre- to post-FMT, including zero change. All models used the same splits and training seed 0. **Figure S3c** summarizes AUROC by the unweighted ten-split mean and sample SD. Repeated test partitions overlapped; no independent-replicate significance test was applied.

MGM2 used the frozen paired-CLS classifier with batch size 32, learning rate 10^−4^, weight decay 0.01 and at most 80 epochs. Its checkpoint minimized validation loss. MOZAIC-repro used microbiome-only DenseNet1D with batch size 16, learning rate 3 × 10^−4^ and 30 epochs, selecting maximum validation AUROC. Shared donors did not merge recipient-linked groups. All models used actual training seed 0 for each split.

### Drug perturbation and recovery data preparation

Drug perturbation and recovery used experiments Exp06 and Exp02 from Shi et al. [33], respectively. We independently reprocessed paired amplicon reads rather than using the original curated MAT abundance table or an earlier OTU99 table. Exp06 contained eight SICs, 90 drug labels, 728 treated wells and 16 DMSO controls. Two plate positions shared the Clomifene label, producing one additional well per SIC; both were retained. Masitinib was also retained. The resulting panel contained all 720 SIC–drug combinations.

Perturbation reads were processed with DADA2 1.30.0 [46] using truncation lengths 240/180, maximum expected errors 2/2, no ambiguous bases, truncation quality 2 and PhiX removal. Samples were denoised separately, reads merged with at least 12 bases of overlap and no mismatches, and chimeras removed by consensus. Error learning used 10⁸ bases, seed 42 and eight threads. All 744 samples were retained, yielding 521 ASVs. Each DMSO well was closed to relative abundance before the two control compositions per SIC were averaged equally. This concurrent control was the reference composition, not a longitudinal pretreatment sample from a treated well.

The perturbation output vocabulary comprised the union of ASVs reaching relative abundance 10^−4^ in any of the eight supplied control states: 163 ASVs plus OTHER. OTHER aggregated remaining mass and was retained in reconstruction and scoring. Vocabulary construction used control compositions only; all eight query communities were known to the predictor.

Recovery used SIC0 exposed to 23 initial drugs or DMSO, with two reseeding arms and two replicates. These 96 trajectories supplied 288 measurements across R1–R3. A separate DADA2 run used truncation 240/180, maximum expected errors 2/2, overlap 12, no merge mismatches and consensus chimera removal. All three passages were denoised together without choosing settings from predictive performance. The input-only R1 vocabulary contained 117 ASVs plus OTHER; held-out R1 inputs could contribute, but R2/R3 outcomes could not. ASV sequences were annotated separately for the two tasks by exact matches to the author-sequence mapping. Ambiguous or absent genus labels remained Unclassified, with family-level fallback when supported.

### Frozen microbial and drug representations

MGM2-Small, Medium, Large and XLarge supplied frozen ASV-token and CLS states of dimensions 224, 320, 384 and 512. ASV nucleotide sequences were encoded by the existing NTv3-650M pipeline before MGM2 contextualization; these sequence embeddings were not an additional downstream feature branch. Drug inputs used DMSO control states, whereas recovery inputs used measured R1 states. Only input ASVs with relative abundance at least 10^−4^ formed local queries; OTHER was excluded. The MGM2 encoders were not fine-tuned.

Drug representations used the frozen Uni-Mol v1 all-hydrogen checkpoint [34] mol_pre_all_h_220816.pt. Each drug had one seed-42 RDKit conformer with hydrogen atoms retained. The cache aligned drug keys, SMILES, atom symbols and conformer coordinates. All valid atom representations had 512 coordinates, with 15–154 atoms per drug and no truncation or planar conformer fallback. Padding and special tokens were excluded. The global drug input was the masked arithmetic mean of valid atom states, not Uni-Mol CLS; the local branch used full atom states, including hydrogens.

### Drug inputs and representation ablations

For each MGM2 scale, the global community vector combined the full CLS state with an abundance-weighted mean of active ASV states. Token weights were control abundances normalized over active queries. The concatenation was standardized using fitting input states, with standard-deviation floor 10^−6^, before appending the unchanged 164-column raw-abundance vector. Its dimension was therefore 164 + 2D, with no global PCA. Local ASV states underwent full-SVD PCA32 fitted on active tokens in fitting control communities. Scores were divided by fitting-population component standard deviations, floored at 10^−6^, and inactive rows were zeroed. No treated outcome entered these transformations.

Uni-Mol atom standardization gave fitting drugs equal weight and valid atoms equal weight within each drug; the global atom-mean vector was standardized separately across fitting drugs. Inner-fold transformations used only inner-training inputs, and final transformations were refitted on the 72 outer-training drugs. The raw-abundance comparator used the unchanged fixed table globally and scalar abundance queries locally, with the same Uni-Mol inputs and predictor family.

### Drug prediction head and model selection

The global branch concatenated community context with the 512-dimensional drug atom mean and used two width-256 linear/SiLU/dropout layers followed by 164 outputs. The local branch projected ASV coordinates and global community context to a shared width, summed them and applied SiLU. Atom states were projected to the same width with SiLU and LayerNorm. Four scaled-dot-product attention heads, with padding masked, were followed by an output projection and residual query update. A shared LayerNorm/linear/SiLU/dropout/linear readout produced one correction per active ASV; inactive ASVs received zero local correction. The initial local width was 128, with widths 32 and 64 additionally evaluated for XLarge.

Global outputs were added to local corrections multiplied by a learned sigmoid gate. The gate logit started at −2, and final global and local output layers started at zero. We divided the combined response by eight, applied tanh and multiplied by eight to obtain a bounded log-compositional response. Adding this response to the log of the reference composition plus 10^−6^, then applying softmax, gave the uncalibrated prediction. Response calibration geometrically interpolated reference and predicted compositions at scale s and reclosed the result. Scale one returned the prediction directly; scale zero returned the reference exactly. Intermediate scales used a 10^−6^ reference pseudocount and prediction floor 10^−30^.

Drug labels defined five folds of 18, with all SICs and duplicate wells for a drug kept together. Each fold served once as the outer test set. Within the remaining 72 drugs, four inner folds used 54 drugs for fitting and 18 for validation. Mean inner-validation Bray-Curtis dissimilarity selected one configuration per model and outer fold, which was then refitted on all 72 non-test drugs before predicting the 18 test drugs. The combined outer predictions covered all 90 drugs, 720 SIC–drug conditions and 728 treated wells exactly once. This design evaluated unseen drugs in the same eight known communities. The random seed for search and final fitting was 42.

Training minimized condition-balanced per-well Bray-Curtis dissimilarity using AdamW with gradient-norm clipping at five. Each epoch used one accumulated update over chunks of at most 512 wells. The initial grid combined learning rates 0.001/0.003, dropout 0.1/0.3, weight decay 0.01, checkpoints at epochs 50/150/400 and response scales 0.5/0.75/1, plus exact no effect, yielding 37 candidates per model and outer fold. For XLarge, a subsequent search compared local widths 32, 64 and 128 at weight decay 0.01, plus width 128 at weight decay 0.1. The global MLP width and frozen inputs were unchanged. Applying the same grid to these profiles yielded 145 distinct candidates after deduplicating no effect. Four-inner-fold mean Bray-Curtis dissimilarity selected the profile and configuration; ties followed the saved profile and candidate order. Selected widths for outer folds 0–4 were 64, 32, 64, 128 and 128, with weight decay 0.1 in the last fold. Other representations retained 37 candidates.

The capacity search reused an already explored dataset; the nested protocol separated fitting and selection within the current evaluation, with different search budgets for XLarge and the other representations. All reported XLarge scores, response examples, attention maps and masking analyses used the same fold-specific inner-selected pipeline. Saved transforms and checkpoints were checked against the corresponding outer predictions before intervention. Additional fixed-checkpoint diagnostics replaced attention by uniform weights or disabled the local branch. Uniform attention gave Bray-Curtis dissimilarity 0.268723 and response cosine 0.442683, compared with 0.270339 and 0.430196 for learned attention; the displayed capacity analysis is shown in **Supplementary Fig. 4b**.

### Drug reference predictions and scoring

Context mean averaged post-drug compositions over the 72 outer-training drugs in the query SIC, first averaging duplicate wells within each SIC–drug pair and then weighting drugs equally. Perturb mean averaged the held-out drug’s measured composition across the seven other SICs after duplicate averaging, excluding every target-SIC well. It is an oracle-information reference because it uses outcomes unavailable for an entirely unmeasured drug.

Bray-Curtis dissimilarity was half the L1 distance between closed predicted and observed profiles. Response cosine compared predicted-minus-reference with observed-minus-reference vectors, assigning zero when the product of norms was at most 10^−12^. The reference was DMSO for drug prediction and measured R1 for recovery. OTHER participated in both metrics. Drug scores were computed per treated well, averaged within SIC–drug pairs, then equally across SICs and drugs. Outcomes were not averaged before nonlinear scoring, and weak-response or poorly predicted conditions were not removed.

### Recovery prediction and nested evaluation

Separate tasks predicted R2 and R3 directly from measured R1; R3 was not predicted recursively through an estimated R2. The 24 initial conditions, comprising 23 drugs and DMSO, were assigned to five folds using seed 20260911. All reseeding arms, replicates and time points of a condition remained together. Each outer fold used the remaining four condition groups for inner validation and then all non-test conditions as response references. Each horizon had 96 scored trajectories. Drug labels were used for grouping only; neither drug identity nor Uni-Mol features entered the recovery predictor.

The decoder transferred same-ASV responses between R1 states within the same known reseeding arm. MGM2 supplied full local ASV states and global CLS without PCA. The abundance comparator used local log1p(b/10^−4^), where b is the ASV relative abundance, and global square-root composition. Candidate geometries were raw coordinates, per-vector L2 normalization, and fitting-input centering followed by L2 normalization. Local squared Euclidean distances were calculated separately for each ASV and scaled by the median positive fitting-reference distance. Global distances were similarly scaled over fitting states. Only input-present ASVs with abundance at least 10^−4^, excluding OTHER, were eligible local references.

For each query and ASV, eligible same-arm references received weights proportional to the exponential of the negative local/global distance mixture divided by a bandwidth. Maximum eligible log weight was subtracted before exponentiation, and weights were normalized over eligible references. Missing query tokens, absent same-ASV references and OTHER used the fitting-mean response in the matching arm. The stable response-transfer implementation was used for every reported result.

Transferred responses used raw differences, square-root differences or log ratios with pseudocount 10^−6^ clipped to [−8, 8]. Responses were scaled and reconstructed in the matching space, negative abundances clipped to zero, and compositions closed. Zero-total predictions reverted to the input composition. The shared grid crossed three geometries, local-distance mixture weights 0/0.5/1, bandwidths 0.03125/0.125/0.25/1/4, three response spaces and scales 0/0.5/0.75/1/1.5/2. Its 810 transfer specifications, 18 matched-arm mean-response specifications and exact no effect gave 829 candidates per representation. Condition-balanced inner Bray-Curtis dissimilarity selected the outer-fold configuration.

Recovery scores averaged replicates within arm, arms within initial condition and the 24 conditions equally. **Figure 4g** reports condition-balanced means from the outer evaluation. For the Rolitetracycline illustration, eligible taxa required a supported genus, mean absolute observed response of at least one percentage point and correct predicted direction in all four arm-by-horizon means. The three largest observed responses were displayed. Lines average replicates within arm and join directly predicted R2/R3 states to measured R1.

### Community attention and atom token interventions

Attention analyses restored the selected MGM2-XLarge outer checkpoint for each test drug and checked its predictions against stored outputs. The input mask retained 58, 43, 51, 35, 52, 30, 41 and 42 ASVs in SIC1–SIC8, respectively: 352 SIC–ASV states and 163 unique ASVs. All 90 drugs therefore supplied 31,680 drug–SIC–ASV queries. For each drug, attention was averaged over four heads, weighted by normalized control abundance within each SIC, restricted to heavy atoms and renormalized. The eight SIC distributions were then averaged equally. Multiplication by heavy-atom count made one denote uniform attention. All displayed molecules shared a single scale derived from their common maximum. Aggregation weights used control abundance only.

Each active ASV’s heavy-atom attention ranking was correlated with the equal-ASV mean of all other active queries in the same SIC and drug. The empirical cumulative distribution gave equal weight to drugs, SICs within drugs and ASVs within SICs. Undefined correlations were excluded from the finite-correlation distribution. We separately compared abundance-weighted and equal-ASV community rankings across all 90 drugs. Illustrative drugs were ranked by outer-test response cosine, with alphabetical tie-breaking. Clotrimazole ranked first and supplied the response example; its eight displayed taxa had supported genus or family annotations and the largest mean absolute observed changes across SICs. The six highest-ranked drugs supplied the molecular attention maps. Both selections were post hoc illustrations and did not use masking effects.

For every held-out drug and SIC, we excluded the top, bottom or a random 10%, 20% or 30% of heavy-atom tokens from all active ASV queries, rounding counts upward and renormalizing attention. Random controls used ten equal-sized masks with generator seed 420000 + 1000 × drug index + the integer percentage masked. Hydrogen tokens, frozen Uni-Mol encoding, global drug input, parameters and calibration were unchanged. The intervention excluded local decoder tokens without physically deleting atoms or re-encoding molecules.

Prediction shift was Bray-Curtis distance between original and masked predictions; accuracy change was masked-minus-original Bray-Curtis error against measured wells; response-cosine loss was original minus masked cosine. We scored individual wells, averaged duplicates within SIC–drug pairs and random-mask scores over ten masks, then weighted SICs and drugs equally. The export contained 25,920 intervention records. **Figure 4f** and **Supplementary Fig. 4d** display 30% masking, selected after examining the 10%, 20% and 30% analyses. Descriptive 95% percentile intervals used 2,000 bootstrap resamples of the 90 drug-level records with seed 420987. The paired high-minus-random Bray-Curtis error interval excluded zero at 30% but included zero at 10% and 20%. These fixed-model intervals did not adjust for selecting the display fraction, historical model exploration or dependence from overlapping training folds. Drug-level summaries retained both positive and negative differences.

Drug-space visualization projected the atom means of all 90 drugs by full-SVD PCA30 followed by two-dimensional t-SNE, with perplexity 20, PCA initialization, automatic learning rate, seed 42 and 1,500 iterations. This display transform was separate from local PCA32 in the predictor. Target labels were matched to source plate metadata by normalized drug names. Pooling unmatched Masitinib into Other yielded 45 human, 32 bacterial, 3 fungal and 10 other labels without assigning a new target. Taxon labels required unambiguous exact ASV-sequence matches; ASV numbers were not transferred between perturbation and recovery vocabularies.

### SAE training on MGM2 OTU-token hidden states

SAEs were trained on MGM2-XLarge OTU-token states from 500,000 microbiome samples, partitioned into 450,000 training, 25,000 validation and 25,000 test samples. The activation dataset contained 124,672,250 OTU-token states and 500,000 CLS states; only the OTU-token states were used for SAE training and interpretation. Training and validation supplied 118,454,598 OTU-token states for fitting, feature-score construction and description generation. The held-out test set was reserved for reconstruction evaluation and did not contribute to feature descriptions.

The primary SAE used BatchTopK with 512-dimensional inputs and a 4,096-feature overcomplete dictionary. BatchTopK retained 64 times the batch size activations across each batch, giving an average L₀ of 64 active features per token. The auxiliary-loss weight was 0.03125. Training used batch size 16,384, learning rate 7 × 10^−5^, optimizer betas of 0.0 and 0.95, no weight decay and 1,000 warmup steps. The model was trained for 61,035 steps with mixed bfloat16 precision. The minimum-validation-loss checkpoint had validation loss 0.032, explained variance 0.96 and mean L₀ of 64.

For SAE training,

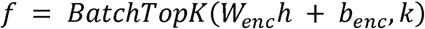

and

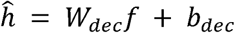

The objective was

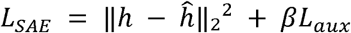

with *BatchTopK* enforcing an average of k active features per token and the auxiliary term encouraging recovery of residual structure from inactive or underused features.

We compared reconstruction and feature usage across BatchTopK settings k = 8, 16, 32 and 64 on 819,200 sampled test tokens (**Supplementary Fig. 5a**). The selected 4,096-feature model used k = 64 and auxiliary-loss weight 0.03125. Training-time deadness was defined by the inactive-counter threshold; global interpretation deadness meant no positive train/validation activation. Inactivity in the sampled test set was a separate quantity. The globally dead features were 3595 and 3795, leaving 4,094 features for interpretation.

### SAE feature evidence construction

We generated evidence packages by streaming train/validation OTU tokens through the selected SAE. For each feature, up to 5,000 high-activation events were retained for interpretation. This cap governed evidence retention; global firing statistics were calculated across the streamed train/validation set. Global summaries included positive count, positive fraction, activation sum and maximum activation.

Evidence packages linked activation events to taxonomy, abundance ranks, sample metadata and project annotations. Deterministic analyses summarized recurrent OTUs, lineage-aware taxon ladders, taxonomic and metadata enrichment, abundance associations, co-occurring taxa and artifact diagnostics. Negative controls matched taxonomy, biome, abundance rank or study. Separate diagnostics assessed project concentration, technology bias and sample concentration when assigning feature descriptions.

### Structured interpretation and assessment of sparse features

Codex, using GPT-5.5 with reasoning effort set to high, generated structured feature descriptions from deterministic evidence packages. The workflow comprised observation, grouping, group description, group validation, unification, final validation and classification. Quantitative evidence, including enrichment statistics, was supplied by deterministic code. Prompts required evidence anchors, assessment of alternative explanations and negative controls, artifact checks, confidence scores and interpretive qualifications.

Each feature retained a unified description, primary class, confidence and acceptance judgment. We joined the initial 500-feature catalog with the remaining 3,594 non-dead features by feature identifier, preserving their existing descriptions and assessments. Confidence was taken from final assessment, or from classification when no final value was present. Primary class and acceptance were treated as separate fields. MGM2Atlas provides interactive access to the complete interpreted catalog, evidence summaries and assessment labels at https://www.mgm2atlas.com/.

### Population habitat profiles and study clustered enrichment

Full-validation summaries comprised 6,212,345 OTU-token events from 25,000 samples and 8,959 studies. Habitat totals were 1,624,619 soil, 1,257,518 plant, 898,432 animal, 1,629,977 aquatic and 801,799 missing. For each feature, habitat-origin shares divided its positive events in a habitat by all of its positive events, retaining missing metadata. Enrichment divided this share by the habitat’s share of all validation tokens. In contrast, within-habitat firing divided positive feature events by all tokens in that habitat. The latter distributions included every one of the 4,094 interpreted features and retained zero firing. A feature with no validation activation had an undefined origin share, not a zero share.

Study-level counts were obtained from a frozen evaluation-mode SAE pass using the original validation indices, four contiguous shards and 16,384-token batch boundaries. All 4,096 features participated in BatchTopK gating. Float32 centering, normalization and encoding were retained, without autocast, TF32, gradients or inactive-counter updates. Feature-by-habitat counts exactly reproduced the original counts in each shard. Preserving batch boundaries retained the original gating context.

For a feature and habitat, the tested effect was the feature’s habitat-origin share minus the corresponding validation background share. The one-sided null specified no positive enrichment. Both shares were token-weighted; studies defined the uncertainty clusters. A study’s influence contribution was its observed-minus-expected feature-active count divided by the feature’s total active count, minus its observed-minus-expected habitat token count divided by the total token count. Expectations used the corresponding overall shares and that study’s feature-active or token total. This retained covariance between the feature share and background. The cluster standard error was the square root of the sum of squared study contributions multiplied by G/(G − 1), with G = 8,959.

We used 19,999 null-centered study-cluster Rademacher multiplier draws with seed 20260914. Each study received one random sign per draw, shared across its habitats and features. The observed statistic was the effect divided by the cluster standard error. Each null statistic was the signed sum of study influences divided by the square root of their squared sum. The Monte Carlo P value was one plus the number of null statistics at least as large as observed, divided by 20,000. The test used linearized study-cluster influences with fixed plug-in studentization.

Tests required at least 20 active studies overall, 10 active studies in the focal habitat, 20 token-bearing studies in both the habitat and its complement, and positive standard error. Unsupported tests received P = 1. Benjamini–Yekutieli correction covered all 4,094 interpreted features and four named habitats, totaling 16,376 hypotheses. Missing metadata remained in denominators but was not tested. **Figure 5c** marks positive effects with adjusted P < 0.05. The feature examples were curated after reviewing the complete results and existing annotations, without changing the testing family or threshold. These associations were conditional on frozen representations; validation data had contributed to the original interpretation evidence.

### Selected event summaries and complete sample codes

Detailed taxonomic and host views used each feature’s separate retained top-5,000 positive train/validation events. Event-count shares and activation-mass shares retained the denominator of the complete selected set. Feature 758’s full-validation habitat profile was kept separate from its selected-event taxonomic summary. Its phylum-by-habitat plot displayed four leading phyla, with circle area proportional to activation mass divided by total retained mass. For Streptococcus features 3915 and 412, rank distributions included only Streptococcus events within their respective selected sets: 4,960 and 4,995 events. Host summaries for features 19, 1399 and 3930 retained ruminant, rodent, bee, other-animal, non-animal and missing groups. Unassigned taxa were excluded only when ranking named families, not from the activation-mass denominator.

Complete sample caches contained 84 × 4,096 coefficients for plant/rhizosphere sample DRR177205.DRS093045 and 241 × 4,096 coefficients for WGS dog sample ERR1914191.SRS1873613. Each complete sample formed one frozen BatchTopK inference batch before displayed coordinates were extracted. Token order was within-sample abundance rank, with rank one most abundant. Positions and features were selected using activations and existing annotations. Displayed expansions retained actual zero coefficients and were not restricted to the four largest coordinates. Individual tokens need not have exactly 64 nonzero features because gating acts across the batch. Sparse coefficients describe a decoder reconstruction and are not percentages of variance or downstream predictive attributions.

Study concentration used the separate retained-event sets for features 973, 3930 and 1399. Studies were ranked by activation mass within each set. Top-study share was the largest study’s fraction of total retained mass; N90 was the minimum number of ranked studies needed to reach 90%. Total study support was 2, 84 and 570, respectively, with N90 values of 1, 23 and 270. Feature 973’s second study was retained despite contributing only 0.019% of its mass. Interpretation considered source concentration jointly with WGS technology, dog host and taxonomic composition.

## Data availability

MicrobeAtlas profiles [9] are available at https://microbeatlas.org/ and MGnify data [7] at https://www.ebi.ac.uk/metagenomics/. FMT data from Schmidt et al. [30, 47] are available at https://doi.org/10.5281/zenodo.6611040 (newly sequenced cohorts: PRJEB46777–PRJEB46780). The Su et al. FMT resource [32] is available at https://github.com/ChenSizhe13893461199/FMT_donor_recipient_matching (PRJNA1262688 and PRJNA1263159). Drug perturbation and recovery data from Shi et al. [33] are deposited under PRJNA1297205. The interpreted feature catalog is available at https://www.mgm2atlas.com/.

## Code availability

MGM2 source code, training and evaluation scripts, and documentation are publicly available at https://github.com/HUST-NingKang-Lab/MGM2. MGM2Atlas, the interactive visualization platform for SAE features and their interpretations, is available at https://www.mgm2atlas.com/.

## Author contributions

H.Z. and K.N. conceived the study and designed MGM2. H.Z. developed the model and analysis framework, performed the main experiments, interpreted the results and wrote the manuscript. Y.Z. analyzed the data and performed benchmarking experiments. Y.Q., T.L. and L.Z collected and organized the datasets and metadata.

R.Y. provided computational resources. K.N. supervised the study, secured resources, guided the scientific framing and revised the manuscript. All authors discussed the results and contributed to the final revision.

## Acknowledgements

This work was supported in part by the National Key R&D Program of China (Grant Nos. 2023YFA1800900 and 2018YFC0910502) and the National Natural Science Foundation of China (Grant Nos. 32071465, 31871334 and 81827901).

## Ethics approval and consent to participate

Not applicable.

