## Supplementary material for "MGM2 as a Multimodal Foundation Model for Microbial Community States and Responses": supplentary figures 1,2,3,4,5

### Supplementary Information

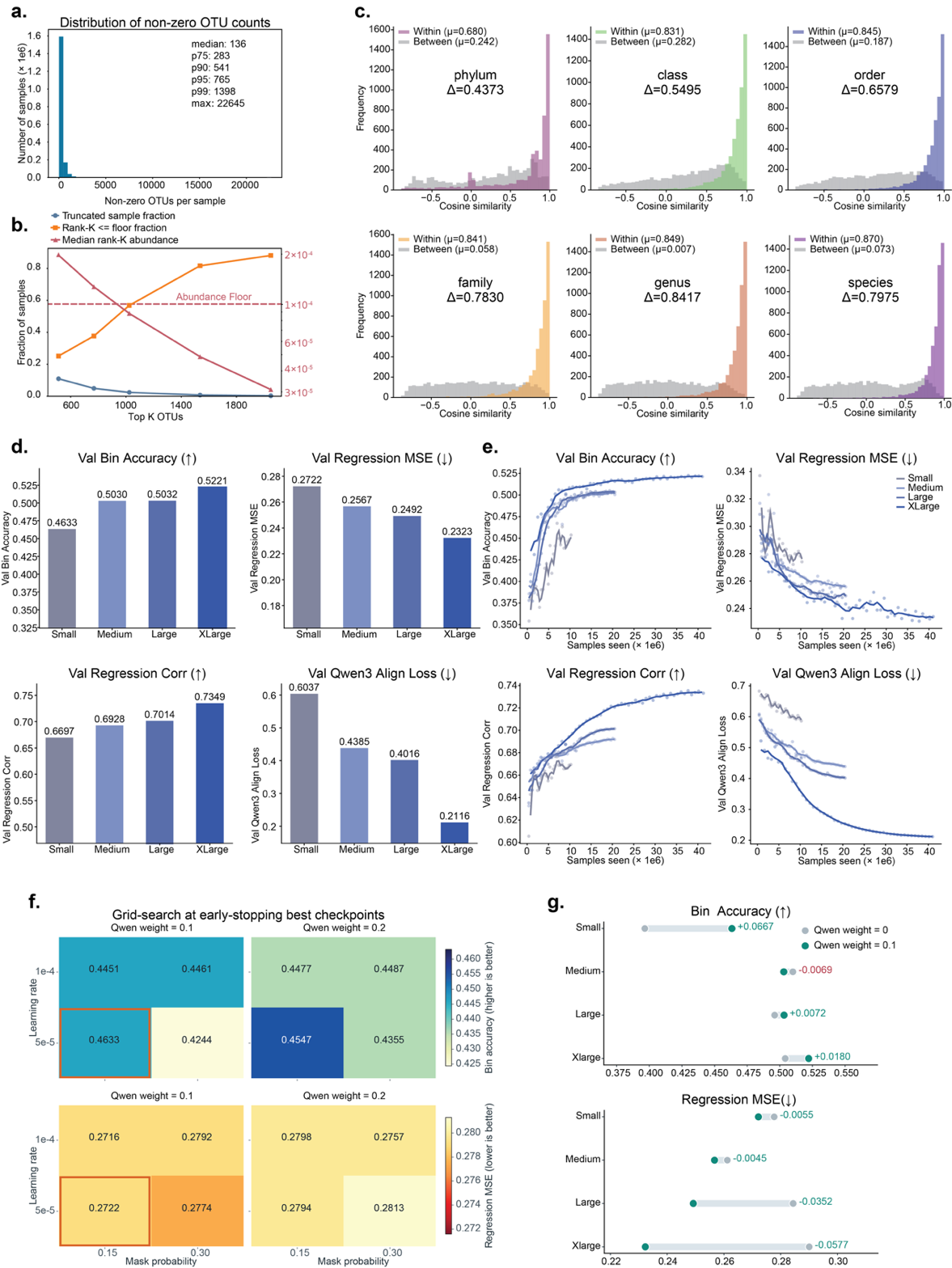

**Figure S1. Pretraining data structure, sequence/semantic ablations and scaling behavior.** **a.** Distribution of non-zero OTU counts across 1,821,291 MicrobeAtlas samples used for MGM2 pretraining. Percentiles summarize the long-tailed complexity of community profiles. **b.** OTU-retention analysis used to set the maximum sample length and abundance floor; increasing the retained top-ranked OTUs reduces truncation while progressively including lower-abundance taxa. **c.** NTv3 representations of OTU nucleotide sequences: embedding similarity within and between taxonomic groups across ranks. Same-taxon pairs show higher cosine similarity than random between-group pairs, with the strongest separation at genus and species levels. **d.** Validation metrics across MGM2 scales, including abundance-bin accuracy, masked abundance regression MSE, regression correlation and Qwen3 semantic-alignment loss. **e.** Training trajectories for the same pretraining objectives. **f.** Early-stopping grid search over learning rate, mask probability and Qwen3 alignment weight; the selected setting balanced abundance classification and regression objectives. **g.** Qwen3 alignment-weight ablation relative to no semantic-alignment loss. Bars show metric deltas for bin accuracy and regression MSE, with lower MSE indicating better masked abundance reconstruction.

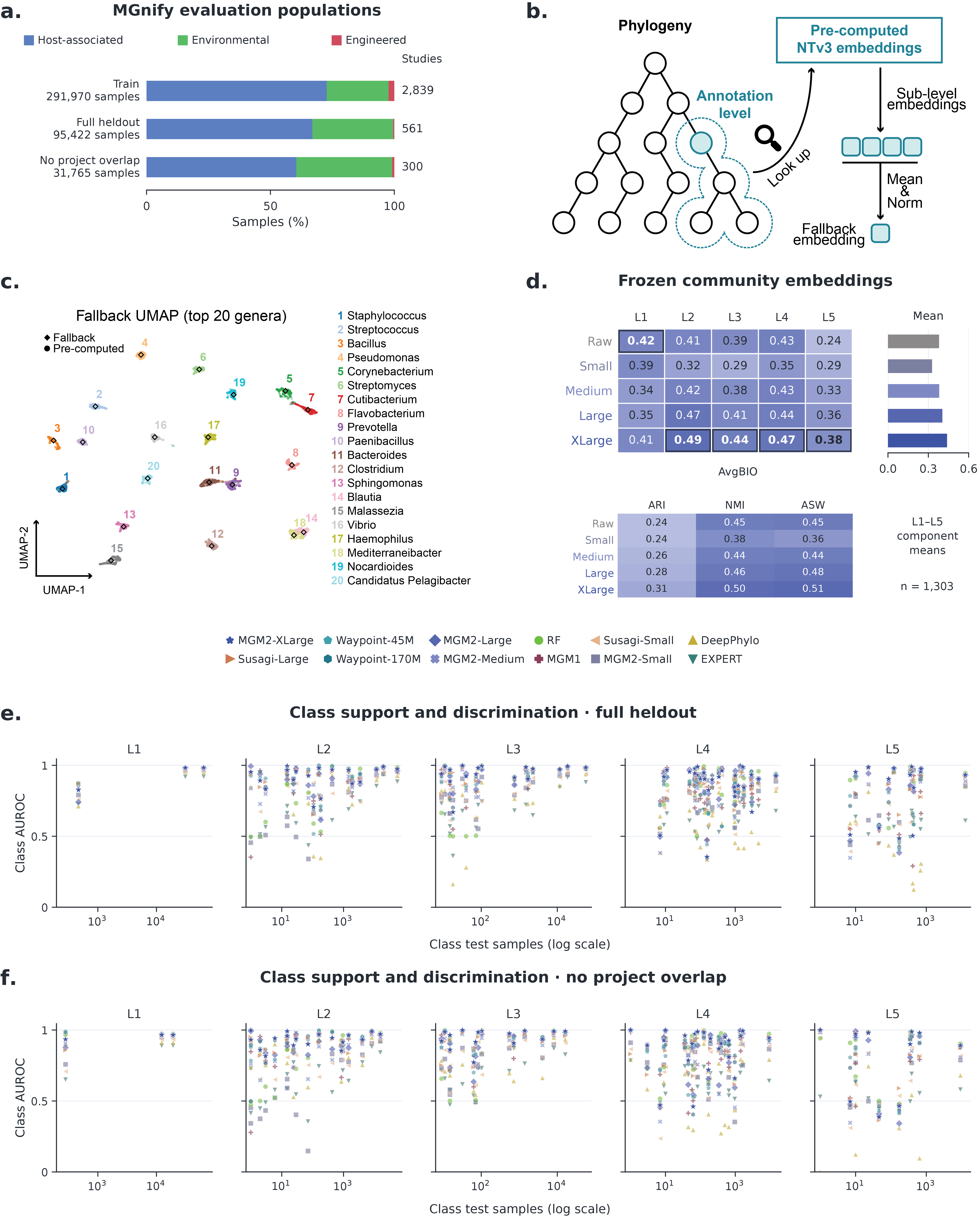

**Figure S2. MGnify evaluation populations and frozen representation structure.** **a.** L1 biome composition of the training, full held-out and project-clean cohorts: 291,970, 95,422 and 31,765 samples from 2,839, 561 and 300 studies, respectively. Project-clean data exclude projects represented in MGM2 pretraining. **b.** Taxonomy-guided fallback for nodes without precomputed sequence-derived embeddings, using the normalized mean of available descendant embeddings. **c.** UMAP comparison of fallback and directly precomputed embeddings for the 20 most frequent genera. **d.** Frozen-embedding organization on 1,303 shared project-clean samples. The upper matrix shows AvgBIO at L1–L5, with the equally weighted five-level mean at right; boxed cells mark column maxima. AvgBIO averages ARI, arithmetic NMI and cosine silhouette rescaled to 0–1. The lower matrix gives the five-level means of these components; ASW denotes the rescaled silhouette component. All representations were L2-normalized, and K-means used 20 initializations with seed 11. Raw abundance used log1p transformation and a 50-component SVD fitted on this subset. Evaluable sample/class counts at L1–L5 were 1,303/3, 1,303/13, 1,132/13, 486/19 and 256/6. e,f. Class support versus mean per-class AUROC for all evaluated classes and 12 models on full (e) and project-clean (f) data. Sample counts use a logarithmic scale. Each point represents a class within a model and hierarchy level. MGM1 denotes the previous MGM framework.

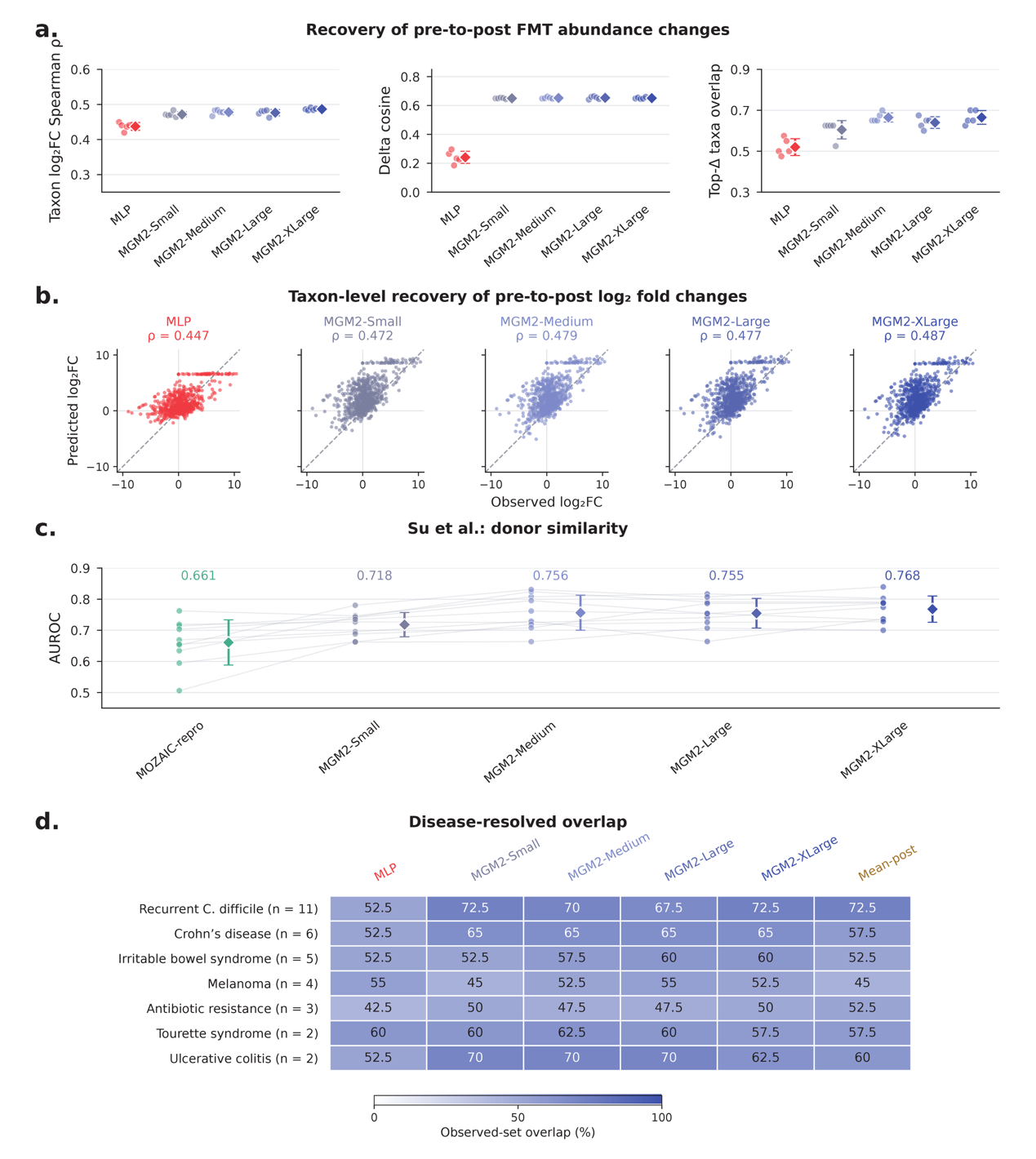

**Figure S3. Recovery of community changes and donor-similarity prediction.** **a.** Change-recovery metrics for the 117-target, 33-patient subset in **Figure 3e**. Circles denote five training seeds; diamonds and error bars show their mean ± sample SD. Spearman correlations compare taxon log₂ fold changes from patient-weighted mean profiles, using a pseudocount of 10⁻⁶ across 755 taxa. Delta cosine compares predicted-minus-recipient with observed-minus-recipient profiles within targets, followed by equal patient weighting. Top-Δ overlap measures recovery of the observed largest positive and negative mean changes, retaining boundary ties. **b.** Observed versus predicted log₂ fold changes across 755 taxa. Predictions are averaged across seeds before patient-weighted profile averaging. Labels report descriptive Spearman correlations of the averaged predictions, rather than the mean per-seed correlations in **a.** Dashed lines denote identity. **c.** Donor-similarity AUROC on the Su et al. dataset (2026; doi:10.1016/j.celrep.2026.117301). The 511 events form 375 recipient-linked groups; ten shared splits assign 270/30/75 groups to training/validation/test. Recipient pre/post samples are disjoint across partitions; groups are defined by sample links, not verified patient identifiers. The positive class denotes non-positive donor-similarity change after FMT, not clinical response. Circles show splits and faint lines connect matched splits; diamonds and error bars show mean ± sample SD. Repeated test partitions overlap, so SD describes split sensitivity. **d.** High-change taxon overlap across all seven indications using the **Figure 3f** selection rule and equal patient weights within each indication. Rows are ordered by patient count and share a 0–100% scale. Observed sets contain 40 taxa; retained ties yield 42, 44 and 42 predicted taxa for Small, Medium and Large in ulcerative colitis. Mean-post remains the fixed training-post baseline. Evaluation details and group sizes are provided in the **main-text Methods**.

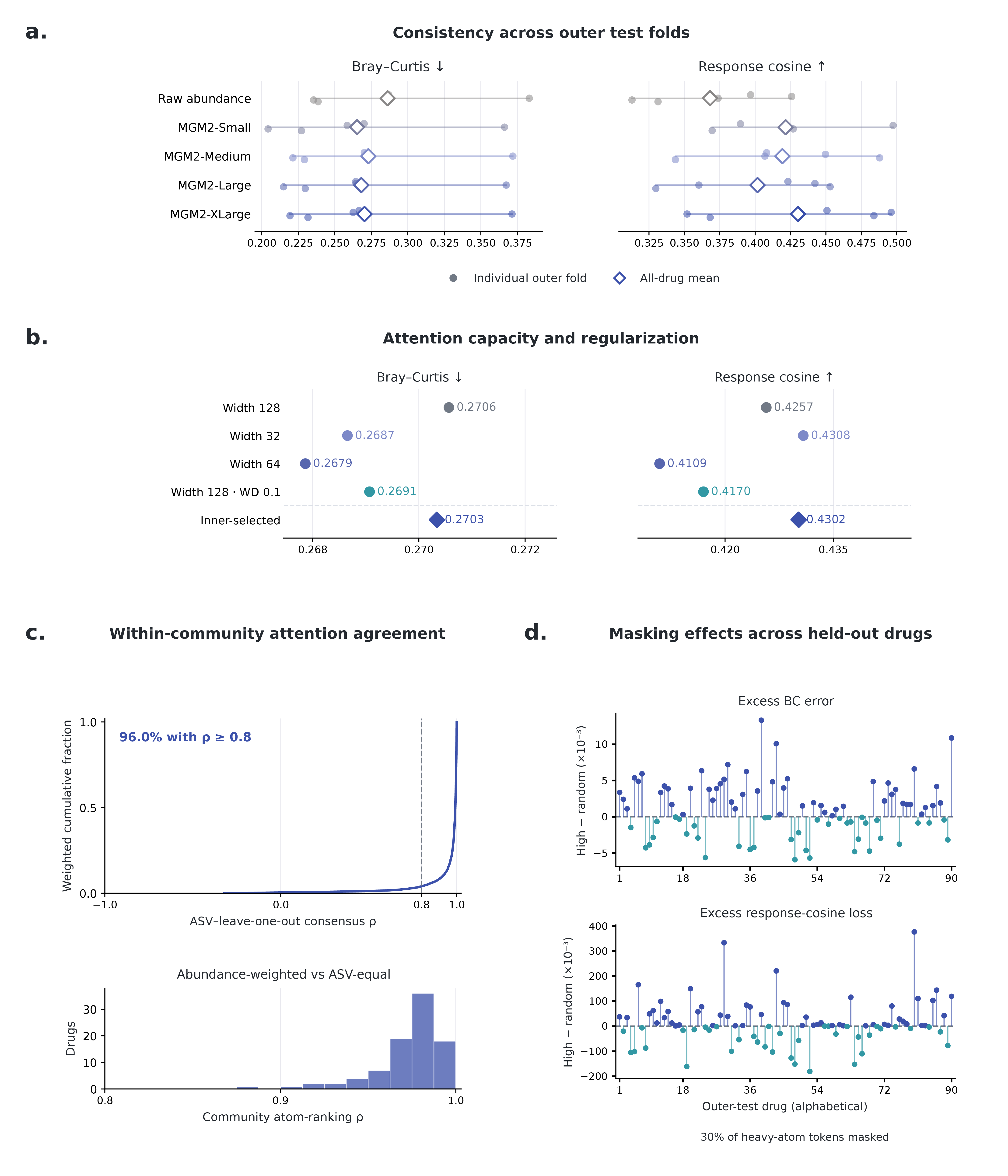

**Figure S4. Five-fold drug prediction and molecular attention analyses.** **a.** Bray-Curtis dissimilarity and response cosine across the five outer test folds for each representation. Circles denote individual folds and diamonds denote all-drug means. **b.** XLarge capacity and regularization profiles: local widths 32, 64 and 128 at weight decay 0.01, width 128 at weight decay 0.1, and the pipeline selecting a profile separately within each outer fold by mean inner-validation Bray-Curtis dissimilarity. **c.** Upper, leave-one-out agreement between each ASV's heavy-atom attention ranking and the mean of the other active ASVs in the same SIC and drug. The finite-correlation distribution weights drugs, SICs and active ASVs equally at each level; 96% have correlation at least 0.8. Lower, abundance-weighted versus equal-ASV community atom rankings across all 90 drugs. **d.** High-attention masking minus the mean random-mask control at 30% heavy-atom masking for all 90 held-out drugs, ordered alphabetically. Upper and lower panels show excess Bray-Curtis error and response-cosine loss, respectively, in 10⁻³ units. Positive values indicate deterioration relative to random masking; negative values indicate improvement. Masking excludes local decoder tokens while leaving molecular encoding and the global drug branch fixed. The displayed fraction was selected from the 10%, 20% and 30% analyses (**main-text Methods**).

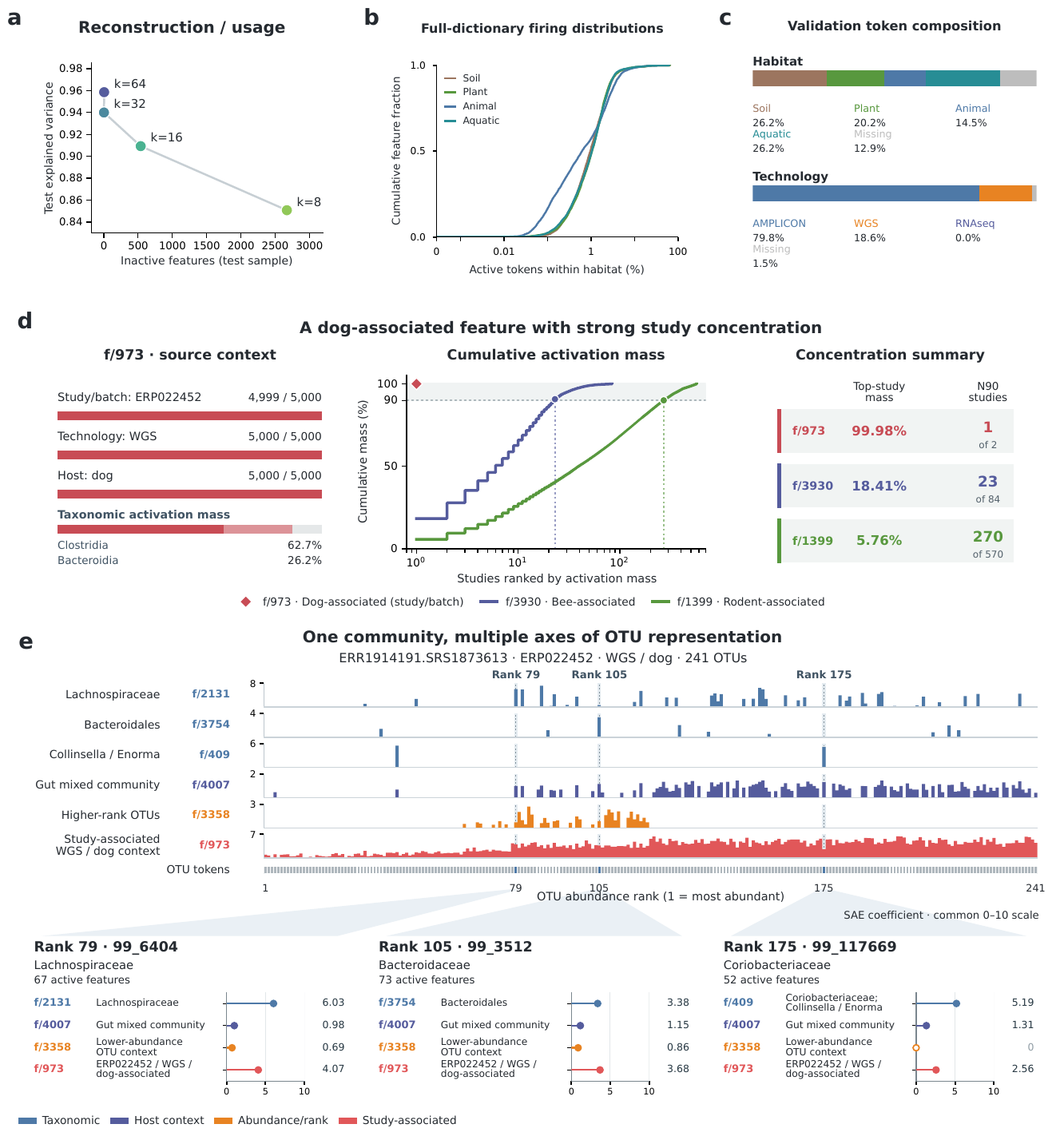

**Figure S5. Reconstruction, population denominators and study-associated features.** **a.** Reconstruction explained variance and inactive-feature counts at k = 8, 16, 32 and 64, evaluated on 819,200 sampled test tokens. **b.** Within-habitat firing distributions over all 4,094 interpreted features, including zeros. Denominators are all validation tokens in each habitat, unlike the activation-origin shares in **Figure 5c,e**. **c.** Habitat and technology composition of 6,212,345 validation tokens, retaining missing metadata. **d.** Feature 973’s retained-event source composition and study activation-mass concentration compared with features 3930 and 1399. Of 5,000 feature-973 events, 4,999 are from ERP022452; all are WGS dog samples. The first feature concentrates nearly all activation mass in its leading study; the other top-study shares are 18% and 5.8%. N90, the minimum number of studies reaching 90% of mass, is 1, 23 and 270 out of 2, 84 and 570 studies. The red diamond retains the unrounded value; the second study remains in the denominator. **e.** Complete 241-token sample ERR1914191.SRS1873613. Feature 973 is active in 235 tokens. Six tracks and selected expansions at ranks 79, 105 and 175 show taxonomy, gut context, rank and study-associated coordinates. Expansions share a 0–10 raw coefficient scale and retain zeros. The selected sample jointly represents source study, host, technology and taxonomic context.
